# On the illusion of scRNA-seq batch effect correction

**DOI:** 10.1101/2025.05.12.653389

**Authors:** Francesco Codicè, Piero Fariselli, Daniele Raimondi

## Abstract

Batch correction methods in single-cell RNA sequencing are essential for removing technical variation that can otherwise lead to misleading downstream analyses. The reliability of these methods is typically evaluated using unsupervised metrics.

Here, we apply a Machine Learning (ML) technique called probing, formalized as the Batch Probing Score (BPS), to empirically demonstrate across six datasets that the most popular batch correctors fail to fully remove batch signal. In most cases, the batch of origin remains clearly identifiable after correction, even though standard evaluation metrics cannot detect it.

We show that existing unsupervised metrics lack the sensitivity and specificity required to capture residual batch signal, whereas ML-based approaches can still detect it. This residual signal can similarly be picked up by downstream analysis tools, potentially leading to biased results.

Because BPS is supervised, it directly quantifies batch signal strength by measuring how accurately the batch of origin can be predicted for each sample. It therefore provides an upper bound on the residual ML-actionable batch signal that could otherwise remain unnoticed.

Our findings suggest that probing-based metrics should become a standard for assessing batch correction methods in single-cell RNA-seq and other areas of genomics.

**Significance statement:** Batch correction is used in virtually every single-cell RNA-sequencing study, yet its effectiveness is often assumed rather than rigorously tested. Removing unwanted technical signal is fundamentally ill-posed, and our results show that widely used correction methods frequently leave substantial batch information behind. Critically, standard evaluation approaches, including visual inspection with UMAP and commonly used quantitative metrics, often fail to detect this residual signal. This means downstream analyses may still exploit technical artifacts and produce biased biological conclusions while appearing properly corrected. By introducing the Batch Probing Score, we provide a more sensitive way to expose this hidden signal. Our study raises an urgent methodological concern for the entire single-cell community and challenges a foundational assumption underlying current analysis pipelines.

## Introduction

Batch correction methods in single-cell RNA sequencing (scRNA-seq) [1, 2] are crucial for removing technical variation (batch effects) that can inflate perceived cell type/status diversity [3], confound downstream analyses [4] and make integrative studies misleading [3]. These batch effects can arise from differences in sample preparation, sequencing runs, reagent lots, or even processing dates [5, 6]. This issue is particularly pronounced when integrating datasets generated with different sequencing technologies, protocols, or in different labs [5, 6]. The goal of batch correction is to remove these confounders by *aligning* the data across batches while preserving genuine biological differences. Distinguishing technical artifacts from true biological variation is a fundamental challenge in life sciences [7, 8, 9, 3], which prompted the development of numerous batch correction methods for scRNA-seq [6, 10, 5, 11, 12, 13, 14, 15].

The first generation of batch correctors, such as ComBat, were designed for bulk microarray data [10, 6], and have later been adapted to scRNA-seq. More recent methods adopted advanced strategies, such as MNN [5] and Harmony [11], which rely on geometric and statistical principles. In parallel, several Deep Learning (DL) approaches address batch effects by learning a shared low-dimensional latent representation across batches, typically with autoencoder-style models. These include methods that use reconstruction objectives and regularization to reduce batch information, generative latent-variable models that treat batch as an explicit covariate, semi-supervised variants that can take into account cell-type labels, and multi-domain translation approaches that align batches through shared latents and consistency constraints [12, 14, 16, 17, 18]. Graph-based methods, like Scanorama [13], construct batch-aware nearest-neighbor graphs to integrate data while trying to preserve the local structure.

From a conceptual perspective, batch correctors should be the *dam* preventing the batch effects accumulated during data generation from *flooding* downstream analyses, and they therefore play a crucial role in safeguarding the meaningfulness and the reproducibility of the results produced by scRNA-seq. But how *watertight* is this dam really? Given their critical safeguard role, scrutinizing their actual efficacy in removing batch effects should be the main priority of the scRNA-seq community.

In practice, in literature, their evaluation relies on qualitative and quantitative unsupervised approaches [5, 13, 11, 14, 6] that range from qualitative, low-dimensional visual inspection (i.e. PCA, t-SNE, UMAP plots), to metrics with an arguably insufficient sensitivity for such a crucial task, such as the Adjusted Rand Index (ARI) and the Average Silhouette Width (ASW) [5, 13, 14, 6], with very few exceptions to this trend [19, 11].

In the qualitative case, dimensionality reduction plots generated with PCA, t-SNE and UMAP are used to visualize the extent of batch signal removal by plotting the data points distribution before and after correction, showing an increased superimposition between data coming from different batches in 2D (see Fig. 1). Unfortunately, reducing the dimensionality of the data from tens of thousands to *two* dimensions means that what we see is just the *2D shadow* of what likely was a very complex distribution of points in an extremely high-dimensional space. Due to the curse of dimensionality, it is therefore hard to assess from these plots whether the samples from different batches can still be trivially separated in the original space, even if they look perfectly superimposed in 2D.

**Figure 1.**
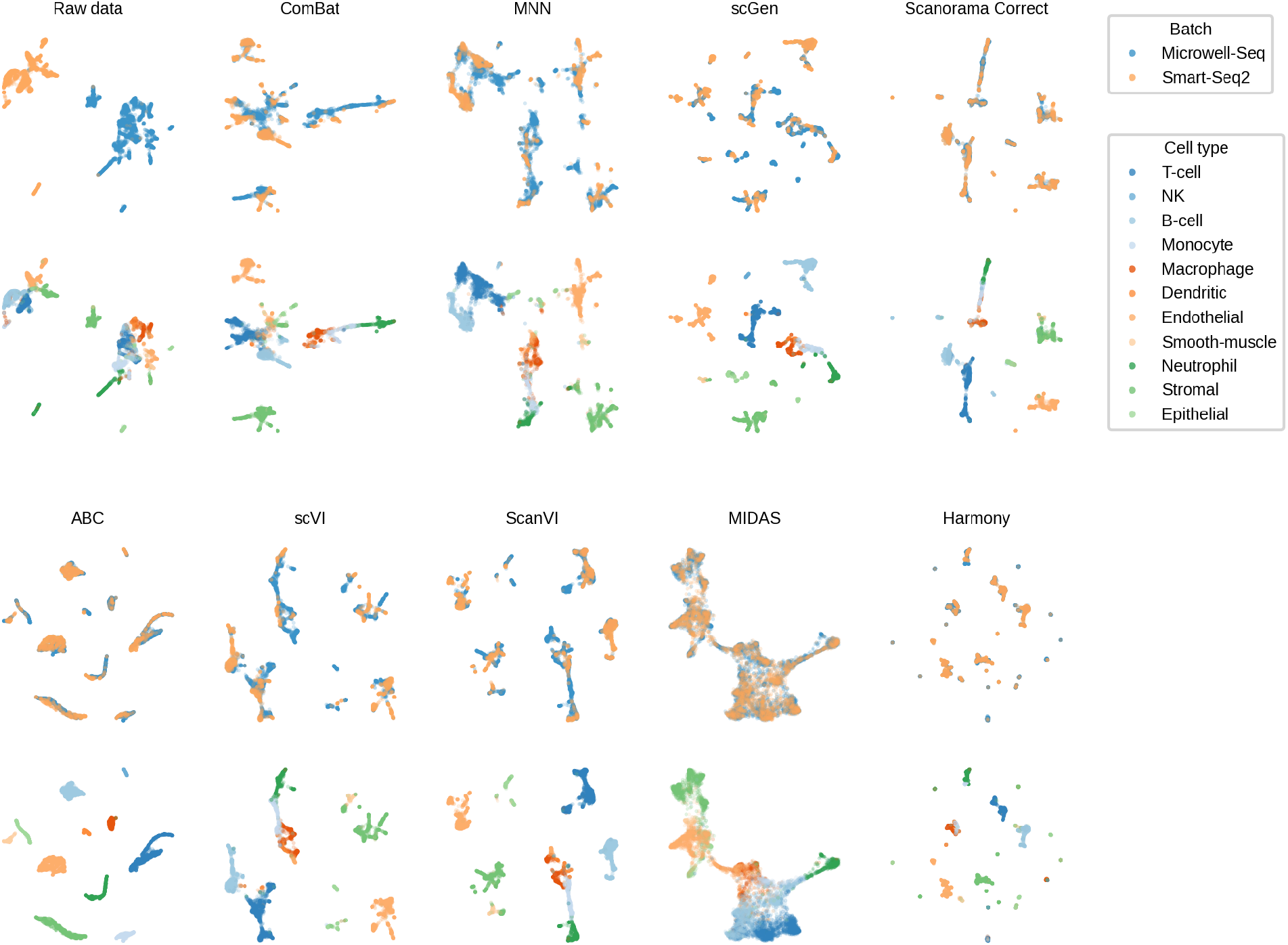
Qualitative overview of batch correction on MMCA2: two-batch scRNA-seq dataset. UMAP projections of the raw data and of the same cells after correction with ComBat, MNN, scGen, Scanorama Correct, ABC, scVI, ScanVI, MIDAS, and Harmony. For each method, the top row is colored by batch (Microwell-Seq vs Smart-Seq2) and the bottom row by cell type, illustrating the trade-off between batch mixing and preservation of biological structure across integration approaches.

Many articles also rely on quantitative evaluations of the residual batch effect after correction [5, 13, 14, 6]. They use unsupervised metrics such as the ARI and ASW, which measure changes in distances between data points and are commonly used to evaluate the goodness of clustering.

Another widely used proxy is PCR (Principal Component Regression), which quantifies how much of the variance in a PCA representation is explained by the batch labels; it is typically computed before and after correction, and the change is interpreted as how much batch-associated variance is removed by the integration step. However, these metrics depend on the chosen embedding and on pipeline choices (e.g., clustering procedure for ARI, and distance definition for ASW), which makes them hard to interpret as objective batch correction scores. As highlighted in [20], ASW is particularly problematic for batch integration benchmarking, because it can be insensitive to structured residual batch effects.

More advanced neighborhood-based metrics, including kBET [19] and iLISI [11], have also been proposed to increase sensitivity to residual batch structure by explicitly assessing local batch mixing. Community efforts such as `openproblems.bio` have collected a broad suite of batch-effect metrics; however, despite differences in formulation and sensitivity, they are all based on geometric or neighborhood-based assessments of batch mixing.

However, even these quantitative metrics share a fundamental limitation: they are *unsupervised*, and therefore rely solely on the spatial arrangement of the data points in the latent space to evaluate the change in batch signal strength. They are not meant to assess whether residual batch signals remain *detectable* or *exploitable* by downstream data analysis tools, and indeed they do not do that at all.

This is problematic because if the batch signal is somehow still hidden in the data after correction, it can bias downstream analyses, especially those involving advanced ML/DL methods (e.g. [21, 22, 23]), which are well known to be able to extract and amplify even *extremely subtle* spurious signals [24, 25].

In the context of scRNA-seq, this can distort embeddings, misclassify cells, and misrepresent biological heterogeneity or trajectories, poisoning all the downstream biological conclusions and results. For example, it could affect DL-based cell type classifiers [21, 22, 23], which may exploit batch-specific artifacts [24, 25] leading to systematic mispredictions [4, 3]. In time-course studies [26], even weak residual batch effects could bias temporal gene expression trends and compromise regulatory network inference [3].

In this study, we show that unsupervised metrics used to evaluate the quality of batch correction methods are unable to really gauge the batch signal strength that remains in the data under correction, and that can still bias downstream analyses and their results. To empirically demonstrate this, we propose the Batch Probing Score (BPS), a supervised metric that quantifies residual batch signal by measuring how well batches can be identified by two standard ML methods, even after correction. Our results show, across multiple real scRNA-seq benchmark datasets and synthetic benchmarks, that existing metrics used to evaluate batch correctors performance lack sensitivity and specificity, and that the BPS can still detect ML-actionable batch signal even when unsupervised, geometry-based metrics indicate that batch signal appears negligible according to those metrics.

Our findings therefore re-assess the true reliability of batch correction methods to be significantly lower than what was currently thought, due to the general inadequacy of nearly all the metrics used to evaluate them. Among the 9 correction methods we tested on 5 public scRNA-seq datasets, only MIDAS [18] is able to substantially reduce the BPS score.

We also empirically show, in a controlled experiment on synthetic data, how the residual batch signal can influence downstream analyses, such as the detection of differentially expressed genes, and therefore the biological conclusions we can derive from these data.

Due to the widespread use of scRNA-seq technologies in research projects around the world, and the avalanche of results they are producing, we urge the community to revise downward the estimate of how much it can rely on batch correction methods.

At the same time, we advocate for *probing-based* supervised metrics, such as the BPS, to be adopted as the new standard evaluation metric for batch correction and similar tasks.

## 1 Results

### 1.1 Probing as a new metric to evaluate batch correction methods

In this paper, we investigate the reliability of the metrics used to evaluate batch correction methods. To do so, we propose a new, *supervised*, metric that aims at measuring the ML/DL-actionable batch signal remaining *after* the correction. We test 9 popular batch correctors on 5 public scRNA-seq datasets (MMCA2,HP5, HPBMC2, MMHSPC2 and HKD11) and a synthetic one. These datasets present different types of batch effects, representative of different real-world situations. MMCA2, HP5 and HPBMC2 suffer from a technology-induced batch effect, but the same cell types are represented in different batches, which is the optimal case for a batch corrector [5]. In contrast, the two batches in the MMHSPC2 dataset contain multiple subtypes of hematopoietic cells, but not all of them are represented in all batches. In the HKD11 dataset, each batch corresponds to a different patient. Since the biological signal of interest is the case-control comparison between healthy and diabetic subjects rather than patient identity, batch correction here aims to remove patient-specific variation while preserving the disease signal. See Section 3.1 for more details.

To quantify the *true* residual batch signal that remains after correction and that could be unknowingly exploited by ML/DL tools used for downstream analysis, we propose the Batch Probing Score (BPS), a new supervised quantitative approach to evaluate the actual effectiveness of scRNA-seq batch correctors by directly measuring the strength of the ML-*actionable* batch signal that remains in the data after correction. Unlike existing metrics (e.g. ARI, ASW), which assess how batch correction modifies the geometric structure of the feature space, BPS tests whether the corrected data still contains sufficient information for a classifier to reconstruct the original batch labels, which is the most direct way to assess the actual extent of the correction.

The BPS takes inspiration from a ML technique called *probing* [27], where one tests whether a target attribute can be predicted from a given representation using a supervised model. In single-cell genomics, probing-style evaluations have been used, for example, to assess generative models by training a discriminator to distinguish *real* from *generated* cells, with ROC-AUC quantifying their separability [28].

Here, we apply the same principle to batch correction. On each dataset, we train a ML classifier before and after correction, to test whether after correction there is still enough signal to predict the batches from which each data point comes and whether the correction reduced such signal at all. We use a Logistic Regression (LR) and a Random Forest (RF) in a cross-validation setting to probe the presence of linear (BPS-LR) and non-linear (BPS-RF) batch signal. A BPS close to 0 indicates that there is no detectable batch signal, while values approaching 1 suggest a strong residual batch effect (i.e. the batches of origin of each sample can be re-identified with perfect accuracy). The BPS classifier is both trained and evaluated, via cross-validation, on the same version of the data being assessed. For example, to quantify the batch signal remaining *after* correction, the BPS is trained and tested on the corrected data.

See Section 3.4 for details on the implementation.

### 1.2 BPS detects batch effect with higher sensitivity and specificity than conventional metrics

Batch correction evaluation strategies based on low-dimensional visualizations of scRNA-seq data can create the illusion that batch effects are largely fixed after correction. The commonly used UMAP plots, such as the ones shown in Fig. 1, show that most batch correctors transform the data so that (i) points belonging to different batches are superimposed and no longer cluster separately, and (ii) cell type structure is visually preserved.

This visual impression is also supported by the widely used unsupervised scores shown in Fig. 2A-B. Clustering/geometry-based metrics, such as ARI and 1 − PCR, suggest near-complete batch removal after correction (mean ARI ≃ 0.06 and 1 − PCR ≃ 0.13, excluding raw data). In contrast, neighborhood-based tests are less optimistic: kBET and 1 − iLISI remain comparatively higher after correction (mean kBET ≃ 0.68 and 1 − iLISI ≃ 0.67, see Fig. 2C and Suppl. Fig. S3), indicating that some residual batch structure is still detectable when evaluated through local mixing criteria. Note that these unsupervised metrics are not fully specific: neighborhood-based metrics can return non-zero values even under randomized null settings (e.g., shuffled batch labels). We show this explicitly in the randomized null results below.

**Figure 2.**
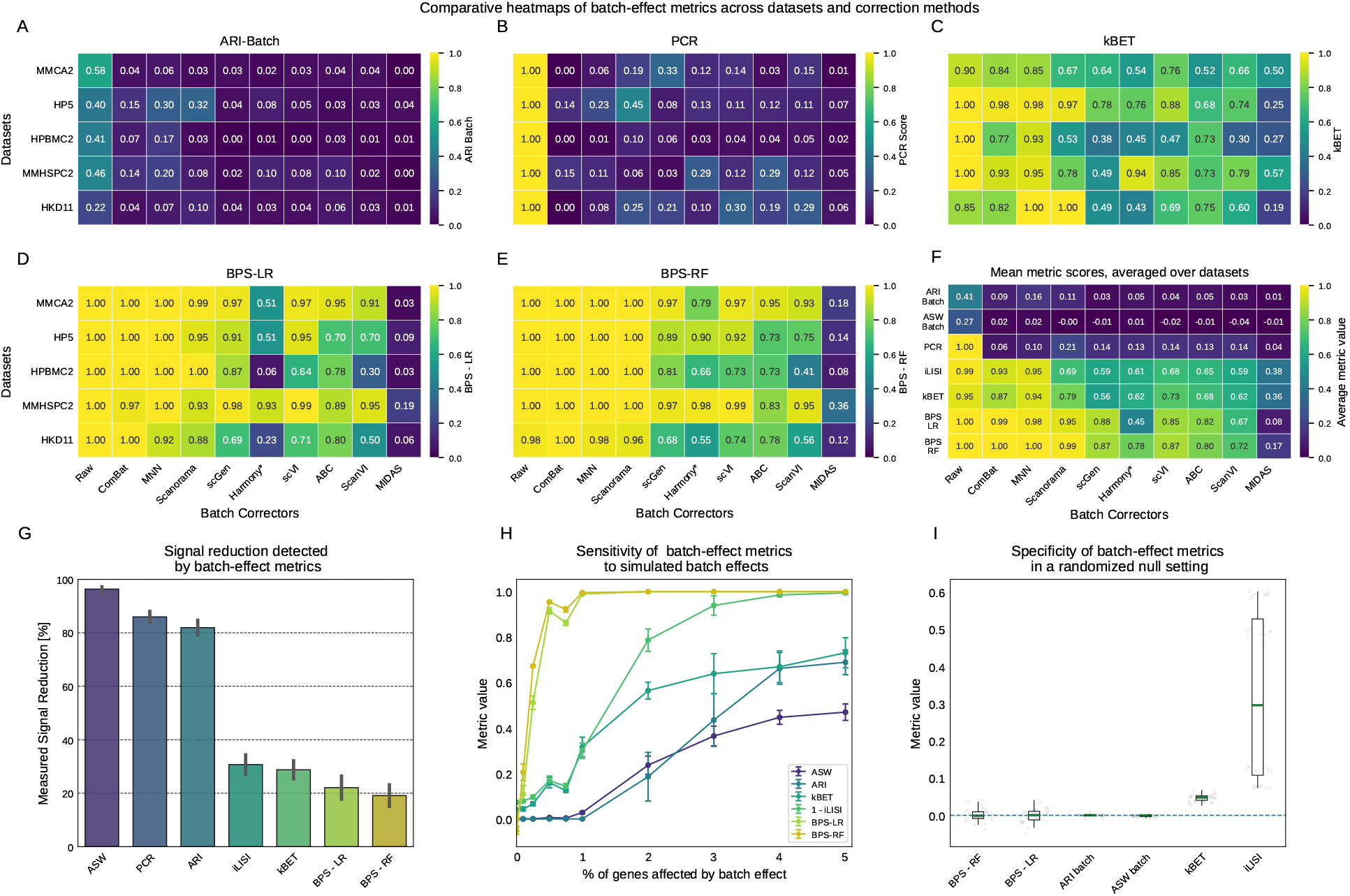
Residual batch-effect assessment across 5 datasets and 9 batch correction methods. (A) Heatmap showing the Adjusted Rand Index (ARI) for batch labels assessing the batch signal strength on 5 datasets (rows) before and after batch correction with 9 methods (columns). (B) Heatmap reporting the principal component regression score (1 − PCR), quantifying the fraction of batch-associated variance explained by the PCA embedding after correction. (C) Heatmap reporting kBET, measuring the fraction of local neighborhoods whose batch composition significantly deviates from the global batch distribution (higher values indicate stronger batch signal). (D) Heatmap showing the same data, evaluated by the Batch Probing Score (BPS-LR). BPS-LR measures the batch signal that can be picked up by a linear predictor (Logistic Regression). (E) Heatmap quantifying the batch signal strength after correction, as perceived by a Random Forest (BPS-RF) non-linear predictor. (F) Summary heatmap (methods ×metrics) reporting mean metric values averaged across datasets, providing a global view of how each batch corrector scores across unsupervised metrics and probing-based scores. (G) Bar plot showing the mean percentage reduction measured by each metric between raw and corrected data, computed across all combinations of batch correctors and datasets (standard error shown). (H) Plot showing the average sensitivity of different batch correction metrics on five simulated datasets (magnitude λ batch = 1). The x-axis indicates the percentage of genes affected by the simulated batch effect (true batch signal strength), and the y-axis shows the average metric values (error bars show variability across simulated datasets). (I) Specificity under a randomized null model: distribution of metric values when batch labels (or batch-associated structure) are randomized, where values close to the baseline indicate that the metric is not spuriously detecting batch signal in the absence of a true effect. *Harmony is the only batch integrator in this benchmark that explicitly operates in a low-dimensional latent embedding (rather than directly correcting gene expression).

However, when we use Logistic Regression (LR) and Random Forest (RF) models to *directly* probe whether batch signal is truly eliminated after correction (see BPS scores Fig. 2D,E), we observe that even a linear model can still reassign samples to their original batches with high accuracy after correction (mean BPS-LR ≃ 0.74, excluding raw data), across multiple datasets and correction methods. Probing residual batch signal with a Random Forest further amplifies this effect: for approximately half of the dataset-corrector pairs, where the batch corrector is not based on complex DL models (e.g. ComBat, MNN, Scanorama, Harmony), batch predictability remains essentially unchanged after correction (BPS-RF ≃ 0.94), while for the remaining more complex methods (e.g. scGen, ABC, scVI, ScanVI, MIDAS), the mean BPS-RF decreases to 0.69. Across all pairs, the mean BPS-RF (excluding raw data) is 0.80, indicating that batch signal is only weakly reduced from the perspective of a non-linear probe. Deep-learning-based correctors reduce batch predictability more than classical methods, but the residual signal is still high. MIDAS is the only clear outlier, since it is able to substantially lower the BPS scores. This is likely due to the fact that this model uses indeed an information bottleneck with two adversarial models used to prevent the batch information from flowing into the final latent representation [18].

Fig. 2E shows that, regardless of what low-dimensional visualization or unsupervised clustering metrics suggest, from the perspective of the ML models on which the BPS score is based, the batch signal largely persists after correction and remains readily exploitable by both linear and non-linear classifiers. Because it directly measures batch-label predictability, BPS provides a more realistic empirical assessment of residual batch signal than geometry-based metrics, which act as weak and easily satisfied proxies.

Consistently, the all-against-all correlations (see Suppl. Fig. S9) show that BPS-derived scores are only weakly to moderately correlated with conventional unsupervised metrics, supporting the view that probing provides non-redundant information about residual batch signal.

Fig. 2F highlights this at the global level: when averaged across datasets, ARI, ASW and 1 − PCR remain close to their no-batch values for most correctors, suggesting near-complete removal, whereas BPS-RF remains markedly higher and better discriminates residual batch signal across methods. Importantly, even the most responsive unsupervised alternatives such as kBET and 1− iLISI stay on average below BPS-RF indicating that probing provides a stricter and more sensitive proxy of batch predictability than both clustering/geometry-based proxies and neighborhood-mixing tests.

This discrepancy is further illustrated in Fig. 2G: ARI, ASW, and PCR report the largest apparent reductions in batch signal, whereas BPS detects substantially smaller reductions. Neighborhood-based metrics (kBET and iLISI) are the only unsupervised alternatives that behave less optimistically and remain more responsive to residual batch structure, approaching the behavior of BPS-LR, which is limited to linear signal. BPS-RF remains more than 1.5-fold more sensitive than the *most sensitive* unsupervised alternative (kBET), making it the most stringent batch-effect metric considered here.

This is also confirmed by the synthetic benchmark in Fig. 2H, in which we injected progressively stronger batch signals into simulated single-cell data (see Methods 3.5 for details). BPS-RF maintains high sensitivity even when the batch signal affects as little as 0.5–1% of genes, whereas the unsupervised alternatives remain unresponsive in these regimes, including kBET [19] and iLISI [11].

Sensitivity alone is insufficient without specificity. In Fig. 2I, we explicitly test the specificity of these methods. To do so, we constructed a randomized null model in which any true batch signal is absent by design: within each dataset we permute ten times batch labels across cells, removing any association between expression and batch, and we then recompute each metric on these randomized instances. Under these null settings, an ideal batch-correction evaluation metric should stay close to zero and not report systematic residual structure. In practice, kBET and iLISI, which are among the most *sensitive* unsupervised metrics,still produce non-zero scores under this setting, showing that they are not fully specific and can be influenced by neighborhood/graph geometry even when batch signal is absent. In particular, iLISI remains above zero under label permutation because its null expectation depends on batch number and batch imbalance, rather than being fixed at zero [11].

In contrast, the BPS remains correctly centered around zero under null conditions, demonstrating the highest sensitivity (Fig. 2H) *and* specificity (Fig. 2I) among all metrics.

### 1.3 BPS provides an upper bound for the statistical power of downstream analysis

To assess how residual batch effects could influence downstream analyses, we designed a synthetic benchmark that allows us to independently control batch-effect strength, biological signal, and alignment between technical (batch) and *“true”* biological signals.

In this synthetic setting, we treat the biological signal as a sample-level property (e.g., case vs control) that induces differential expression on a small set of genes, while the batch signal acts as an independent technical shift affecting a (typically larger) set of genes. We control both the strength of these shifts and the size of the affected gene set. We vary the batch–biology alignment at the sample level: higher alignment means that condition-positive samples are increasingly concentrated in one batch, so batch membership and biological condition become more correlated and mutually predictive.

This synthetic experiment demonstrates, in a controlled way, how performance declines with respect to batch magnitude and/or by confounding with biology.

We select differential-expression (DE) analysis as a representative downstream task: we compare the two simulated biological conditions, we rank genes by DE evidence, and we quantify the recovery of DE genes using the AUPRC against the set of genes perturbed by the synthetic “biological” signal. Fig. 3A shows that the recovery of DE genes progressively degrades as batch effects become stronger and increasingly aligned with biological conditions, showing how confounding and technical signal directly impair downstream performance even when biological signal is present.

**Figure 3.**
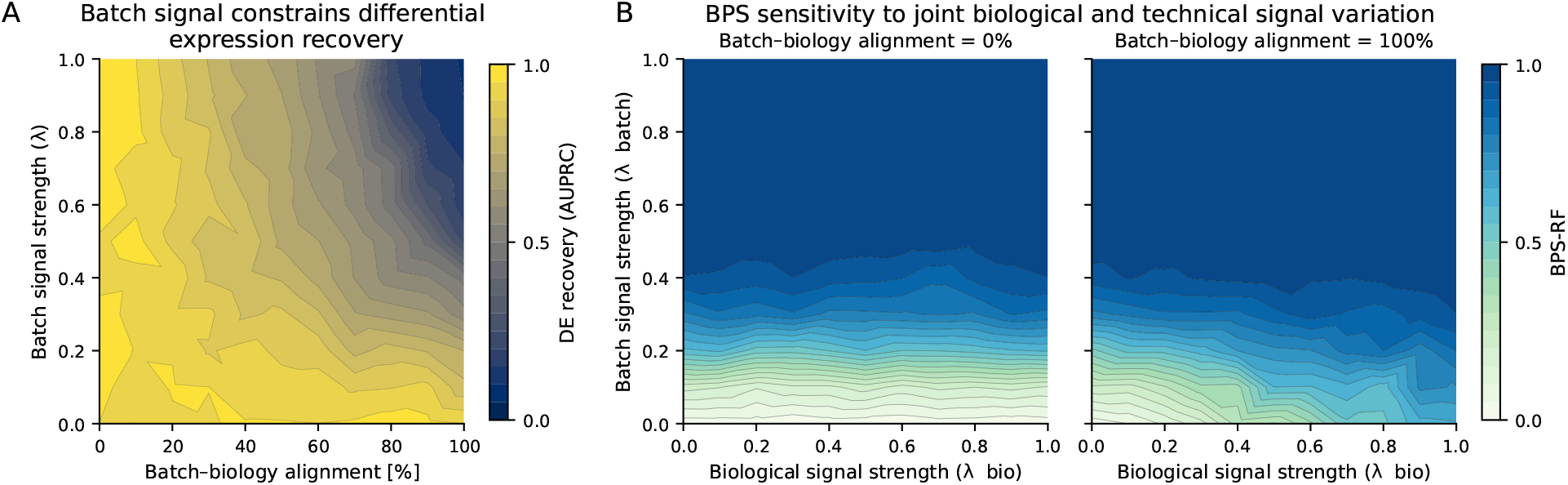
Residual batch signal provides an upper bound on potential batch-driven bias in downstream analyses. (A) Differential-expression recovery in a synthetic benchmark as a function of batch–biology alignment and batch-effect strength, illustrating that increasing confounding and stronger technical signal degrade downstream performance. (B) Batch Probing Score computed with a Random Forest (BPS-RF) across joint variation of biological and technical signal strengths under two levels of batch–condition alignment, showing that BPS increases monotonically with actionable batch signal. Together, these panels support interpreting BPS as a conservative proxy for how much batch information remains exploitable by expressive downstream models.

This emphasizes why we should explicitly quantify residual batch signal after correction: while we illustrate the effect with a simple and interpretable downstream analysis (DE), the same residual signal can be even more problematic in ML/DL pipelines, where batch-driven shortcuts are harder to spot and separate from biology due to model complexity and limited interpretability.

Fig. 3B shows BPS-RF as we vary biological and batch signal strength, both when batch and biological condition are independent and when they are fully aligned. The left panel of Fig. 3B shows how, across all regimes of biological and technical signal variation, the BPS-RF score increases monotonically with the amount of actionable batch signal under zero batch–biology alignment. When batch and biological signals are fully aligned (right panel of Fig. 3B), any signal correlated with the batch is indistinguishable from the biological signal. In this scenario, BPS cannot distinguish between biological and batch signal, because they are aligned.

In summary, these results show that increasing batch signal worsens DE recovery downstream task (Fig. 3A) and that BPS tracks this residual, exploitable batch-associated effect (Fig. 3B). We therefore interpret BPS as a conservative upper bound on how strongly downstream analyses can be biased by batch-driven signal.

### 1.4 BPS specificity and robustness to cell-type composition effects

Batches may differ not only in their technical gene expression profiles but also in their cell-type composition: if *P* (cell type | batch) varies across batches, this compositional imbalance is itself a source of batch-predictable signal that downstream ML models can exploit, potentially biasing any analysis that does not account for it. This is a specific instance of the covariate shift problem [29, 30], well-documented in machine learning contexts [24]. From this perspective, the global BPS correctly captures composition-driven signal as a genuine component of the residual batch effect that remains actionable after correction.

At the same time, it is useful to disentangle this compositional component from the purely technical, within-cell-type batch signal. To be consistent with established practice in the scRNA-seq integration literature [6], the BPS classifier is trained and evaluated exclusively on cells belonging to cell types shared across all batches (see Section 3.4), preventing the classifier from exploiting the absence of cell types in some batches as a trivial predictive shortcut.

To go one step further in removing the compositional component from the BPS evaluation, we additionally computed a per-cell-type variant of BPS-RF, in which the classifier is trained and evaluated *separately within each cell type*, and the resulting scores are averaged using cell-type frequencies as weights (see Suppl. Fig. S17).

The values remain broadly consistent with the global BPS-RF reported in Fig. 2E, confirming that residual batch signal persists even once the contribution of cell-type composition is fully eliminated.

We further assessed the specificity of BPS using two null models (Suppl. Fig. S16). A random-permutation null model, which removes all associations between cells and batch labels, and a stratified cell-type-preserving null model, which additionally controls for compositional imbalance by permuting batch labels within each cell type, both yielded near-zero per-cell-type BPS values (Suppl. Fig. S16). Under the stratified null model, the global BPS remains non-zero, whereas the per-cell-type BPS remains close to zero: even when batch labels are permuted within each cell type, removing any technical signal, the global score retains sensitivity to composition-driven batch predictability that persists by construction. The per-cell-type BPS, being evaluated within the stratified null, is blind to this component and therefore returns to zero under both nulls. This behavior supports using both variants in practice: the global BPS quantifies the total residual batch signal exploitable by downstream models, including compositional imbalance, while the per-cell-type BPS more cleanly isolates the purely technical within-cell-type component.

### 1.5 BPS is an interpretable metric that identifies genes driving residual batch effects

In addition to quantifying the magnitude of residual batch signal with high sensitivity and specificity, BPS natively provides an interpretable framework to identify the genes carrying the batch signal. Both the BPS-LR and RF models’ predictions can be interpreted by inspecting the LR coefficients and the RF feature importance scores, respectively capturing linear and non-linear gene contributions towards the batch prediction. These feature contributions can be extracted in the form of rankings, indicating which genes are the most correlated with the batch signal.

To show the interpretability of BPS on real data, in Fig. 4A, we assess whether the top-ranked genes are enriched for well-known proxy technical gene sets[31, 32] on the HPBMC2 dataset. The *y* axis shows how many of the top-ranked genes driving the prediction belong to mitochondrial (`MT-`) or hemoglobin (`HBA/HBB`) gene sets.

**Figure 4.**
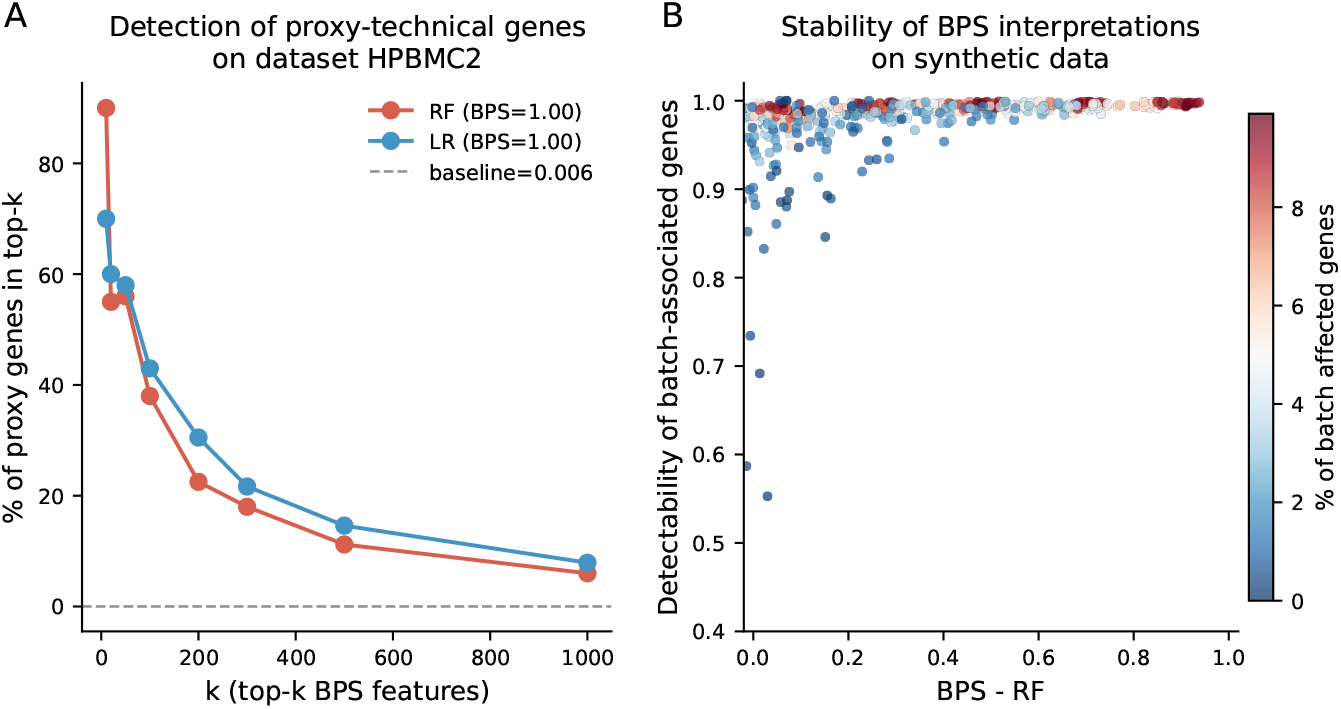
Interpreting BPS through feature attribution on real data and validation on synthetic benchmarks. (A) Enrichment of proxy technical gene sets among the top-*k* most influential features selected by BPS models (Logistic Regression and Random Forest) on HPBMC2; the dashed line indicates the baseline prevalence of the proxy set among all genes. (B) Synthetic benchmark relating residual batch predictability (BPS-RF, x-axis) to the detectability of batch-associated genes (y-axis) at fixed batch-effect strength (λ batch = 0.25); points are colored by the percentage of batch-affected genes.

Consistently, the most influential genes according to BPS are significantly enriched for these proxy technical gene sets, indicating that residual batch signal is primarily supported by technical rather than biological features. Fig. 4A shows that this enrichment is strongest among the top-ranked features and gradually decreases as larger feature sets are considered, yet it consistently remains above the baseline prevalence of technical genes across all values of *k*, suggesting that residual batch effects are not only concentrated in a limited subset of technical drivers.

While the RF is known to be a reliable selector of relevant features[33], also BPS-LR achieves similar results.

Fig. 4B shows, on synthetic data, how the detectability of batch-associated genes (*y* axis) is influenced by the batch effect predictability (*x* axis) and by the percentage of genes affected by the batch effect (color gradient). The results in Fig. 4B indicate that the BPS-RF can correctly retrieve the genes carrying the batch signal consistently, except for regimes with very low batch signal strength and small percentage of affected genes. The interpretability of the BPS score does not require strong residual batch signal to be reliable.

BPS therefore provides a reliable ranking of candidate genes associated with batch, making it straight-forward to identify which genes most strongly drive the residual batch signal.

### 1.6 Joint evaluation of batch removal and biological signal preservation

When it comes to evaluating scRNA-seq correctors, the reduction of the batch signal strength is not the only desideratum. Batch correction is inherently a *min–max* problem: the batch signal must be minimized while the true biological signal should be preserved and allowed to emerge.

In Fig. 5, we evaluate both aspects simultaneously with a 2D plot. On the *y*-axis (ARI–Bio Cluster), we quantify the cell-type signal accessible to unsupervised ARI clustering metric. While ARI is known to be less sensitive than supervised metrics, this lower sensitivity makes it an appropriate and conservative proxy for biological structure preservation. On the *x*-axis, we report the BPS-RF of each method, which captures the residual, ML-actionable batch signal.

**Figure 5.**
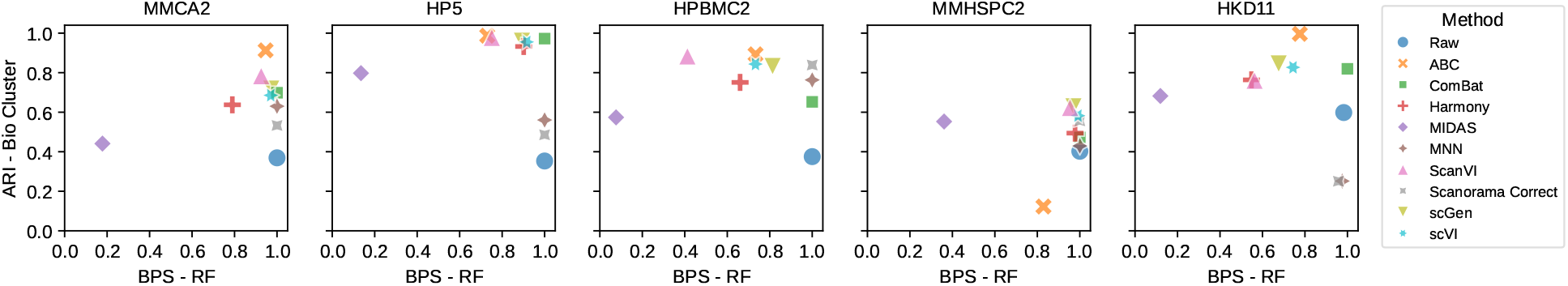
Joint evaluation of batch removal and biological signal preservation. Scatter plots summarizing the performance of batch correction methods across five scRNA-seq datasets (MMCA2, HP5, HPBMC2, MMHSPC2, HKD11). Each point represents one correction method. The x-axis reports the Batch Probing Score computed with Random Forest (BPS-RF), quantifying the residual ML-actionable batch signal (lower is better). The y-axis reports the Adjusted Rand Index (ARI-Bio Cluster) for cell-type clustering, measuring biological signal preservation (higher is better). The ideal correction method would lie in the top-left corner, corresponding to strong preservation of biological structure and minimal residual batch signal.

In this two-dimensional evaluation, the ideal batch corrector would lie in the top-left corner, corresponding to a strong preservation of cell-type structure (biological signal, high ARI-Bio Cluster) and effective removal of batch effects (low BPS).

Across datasets, most methods remain positioned toward the right side of the plot, indicating substantial residual batch signal despite acceptable clustering of the cell types. Among the correctors benchmarked in this study, MIDAS represents a relatively balanced solution. It is generally the only one that can noticeably reduce the batch signal, even if this often comes at the cost of a reduction of the biological signal as well, with respect to the other methods.

Fig. 5 highlights that no single metric is sufficient to evaluate a batch corrector: BPS quantifies how much batch signal remains exploitable, but it should be interpreted jointly with biological preservation metrics such as ARI-Bio Cluster, allowing users to select the method that best suits the specific trade-off required by their downstream analysis.

## 2 Discussion

In this paper, we showed that conventionally used metrics for the evaluation of batch correction methods tend to substantially underestimate the residual batch signal after correction. Low-dimensional visualizations (UMAP/PCA/t-SNE) often suggest near-perfect mixing, while metrics based on clustering or geometry (ARI, ASW, PCR) can report strong apparent batch removal even when batch labels remain highly predictable. Neighborhood-based metrics such as kBET and iLISI are generally more sensitive to residual structure than ARI/ASW/PCR, but they lack specificity: under randomized null settings they can still return non-zero values, indicating spurious detection driven by local graph geometry. Earlier work such as kBET[19] already noted that batch correction is at risk of being over-optimistically assessed with qualitative approaches. Here we go a step further by showing that this holds even for quantitative *unsupervised* metrics, and that a supervised non-linear probe is needed to truly maximise the sensitivity (and specificity) of batch signal detection.

Thanks to its supervised approach, BPS *directly* measures residual batch information as batch-label predictability *P* (*batch* |*data*), and therefore aligns with the question that matters for downstream analyses, such as *“how much of the batch signal remains exploitable by learning algorithms?”* Across real datasets, BPS reveals consistently high residual predictability after correction, and in controlled synthetic benchmarks it achieves the best sensitivity to subtle batch effects while remaining equal to zero when the batch signal is not present, therefore achieving a high specificity.

Our results suggest that most of the current batch correctors leave substantial residual ML-actionable batch signal across datasets. Among the evaluated approaches, DL-based correctors, such as scGen [12], ABC [14], scVI [16], ScanVI [17] and MIDAS [18], tend to achieve better corrections, reducing BPS more consistently than classical statistical/geometric methods such as ComBat [10], MNN [5], Scanorama [13] and Harmony [11], although residual predictability remains non-negligible in most dataset-method pairs. MIDAS is the best corrector, in terms of BPS reduction, but at the price of a modest drop in cell-type clustering performance (lower ARI-Bio Cluster) relative to the best biologically preserving methods. These findings encourage researchers to pursue the development of DL methods for batch correction, and motivate the evaluation of these future approaches with *supervised* batch signal evaluation metrics such as the BPS.

To make BPS more consistent and conservative, we base it on a “One versus the Rest” (OvR) ROC AUC (see Methods 3.4) rather than other classification metrics like the MCC. OvR AUC asks a simpler (and, for our purposes, stricter) question: can a classifier still distinguish one batch *from the others* after correction? The MCC instead emphasizes assigning *each cell to its exact batch of origin*. Since downstream analyses can be biased by the simple fact that batches are *distinguishable*, and not necessarily *re-identifiable*, the OvR AUC is the safer choice (see Section 3.4 for details).

BPS also remains robust to cell-type composition imbalance across batches: the per-cell-type-BPS isolates the within-cell-type residual signal and remains close to zero under a stratified null model, confirming that the residual batch predictability we report is not merely an artifact of differing cell-type proportions (see Section 1.4).

In this sense, our synthetic downstream experiment illustrates the practical consequence of residual batch signal: increasing batch strength or the alignment between the batch and biological signals progressively degrades the recovery of differentially expressed genes (Fig. 3A). In the same setting, BPS increases with residual batch signal (Fig. 3B), tracking the amount of batch-correlated structure that downstream models could exploit. We therefore propose BPS as a conservative upper bound on the potential impact of batch-driven signal on downstream analyses, providing a stringent warning signal when biological conclusions depend on increasingly powerful ML/DL pipelines.

Finally, since it is based on a ML technique called *probing*, that is applicable virtually on any field of science involving the generation and analysis of data, the applications of BPS-like measures are not limited to single cell or scRNA-seq, but could be used to detect batch effects in various types of biological data, including sequencing data, SNP arrays, cell-free DNA, bulk RNA-seq and any other omics.

## 3 Methods

### 3.1 Real-world scRNA-seq datasets

In this section, we describe the datasets used in our study. For each dataset, we report the total number of cells and genes, and specify the type of batch effect being addressed. The benchmark performed on these datasets encompasses various scenarios, including differences in scRNA-seq technologies and batch effects specific to the protocol or patient. This allows us to test the different correction methods in diverse settings. In particular, the MMCA2, HP5, HPBMC2, and MMHSPC2 datasets have been taken already preprocessed from [6]. HKD11 was obtained from the single-cell data integration task of the Open Problems in Single-Cell Analysis benchmark (openproblems.bio). It is adopted as a standard reference for evaluating batch correction methods.

1. **Mouse Cell Atlas (MMCA2)**. The dataset originates from mouse (*Mus musculus*) tissues covering nine organ systems and includes 11 distinct cell types that are consistently represented across batches [34, 35]. It comprises a total of 6,954 cells and 15,006 genes. Data were collected in two batches using different scRNA-seq technologies, with one batch comprising 4,239 cells (Microwell-seq) and the other 2,715 cells (Smart-Seq2) [34, 35]. In MMCA2, our goal is to correct batch effects induced by these technology differences. We followed the pre-processing used in [6].
2. **Human Pancreas (HP5)**. This dataset comprises human pancreatic tissue data representing 15 distinct cell types that are uniformly present across batches [36, 37, 38, 39, 40]. It contains a total of 14,767 cells with 15,558 genes measured per cell and was integrated from five batches profiled using four different scRNA-seq technologies. Here too, the goal is to correct the batch effects caused by technology differences [6].
3. **Human PBMC (HPBMC2)**. The dataset consists of human peripheral blood mononuclear cells (PBMCs) encompassing 9 cell types that are consistently represented across batches [41]. It comprises a total of 15,476 cells with 19,357 genes measured per cell [41]. Data were generated in two batches using different 10x Genomics protocols: one batch with 8,098 cells obtained from the 3’ protocol and another with 7,378 cells from the 5’ protocol. The aim is to correct batch effects resulting from protocol-driven differences [6].
4. **Mouse Hematopoietic Stem and Progenitor Cells (MMHSPC2)**. This dataset contains mouse hematopoietic stem and progenitor cells from various hematopoietic tissues, representing 7 distinct subtypes [42, 43]. It comprises a total of 4,649 cells and 3,180 genes. Data were generated in two batches using different sequencing protocols: one batch with 1,920 cells from SMART-seq2 and another with 2,729 cells from MARS-seq. While both batches contain multiple subtypes, not all subtypes are represented in every batch. Here, we focus on correcting batch effects induced by differences in sequencing protocols [6].
5. **Human Diabetic Kidney Disease (HKD11)**. The dataset originates from human (*Homo sapiens*) renal cortex tissue, focusing on diabetic kidney disease (DKD) [44]. It comprises a total of 39,176 cells and 27,980 genes, collected across 11 batches (6 from healthy subjects and 5 from diabetic patients), and includes 13 distinct cell types [44]. In this dataset, all batches were produced with the same technology, and batch effects are mainly driven by inter-individual variability. HKD11 has been adopted as a benchmark dataset in the data integration task on openproblems.bio. With this dataset, we aim to correct the batch effect among patients to remove technical variability and accurately assess the transcriptional alterations associated with DKD progression.

**Table 1:** Summary statistics of scRNA-seq datasets. For each dataset, gene and cell counts, the number of batches and cell types, and the normalized entropy (representing the uniformity of cell type distribution across batches) are reported. See Suppl. Table S1 for cell-type composition details.

| Dataset | Gene Count | Cell Count | Batch Count | Cell Type Count | Normalized Entropy |
| --- | --- | --- | --- | --- | --- |
| MMCA2[34, 35] | 15006 | 6954 | 2 | 11 | 0.88 |
| HP5[36, 37, 38, 39, 40] | 15558 | 14767 | 5 | 15 | 0.68 |
| HPBMC2[41] | 19357 | 15476 | 2 | 9 | 0.81 |
| MMHSPC2[42, 43] | 3180 | 4649 | 2 | 7 | 0.78 |
| HKD11[44] | 27980 | 39176 | 11 | 13 | 0.83 |

### 3.2 Batch Correctors

We evaluated nine state-of-the-art batch correction methods. This list consists of statistical, geometric, and deep learning approaches. Before correction, gene expression values were normalized by total counts per cell, then log-transformed, and the top 5,000 most variable genes were selected for downstream analysis. Each method aims to remove technical variability while preserving biologically meaningful variation.

Here we provide a summary of the batch correctors we tested:

1. **ComBat**. We applied the ComBat algorithm [10] to address batch effects in single-cell RNA sequencing data, using the implementation provided by the Scanpy toolkit [45]. ComBat uses a parametric empirical Bayes framework to model batch-specific additive and multiplicative parameters, harmonizing technical variation across batches while retaining biologically relevant heterogeneity.
2. **Harmony**. Harmony is an algorithm for integrating single-cell RNA-seq data from multiple datasets. It embeds cells into a low-dimensional space to capture the primary structure of gene expression. Harmony then iteratively performs soft clustering to assign cells to clusters, ensuring that each cluster includes cells from different batches. Within each cluster, batch-specific centroids are calculated and used to derive cell-specific linear correction factors based on the cell’s soft membership across clusters. These correction factors are applied to the original embedding, and the process is repeated until the cluster assignments stabilize. The final output is a corrected embedding in which batch effects are reduced while preserving the biological signal [11].
3. **MNN (Mutual Nearest Neighbors)**. MNN identifies cells of the same type across batches by computing mutual nearest neighbors (MNNs) based on Euclidean distances after cosine normalization. For each MNN pair, a batch-correction vector is derived from expression differences. Cell-specific corrections are calculated as weighted averages of these vectors using a Gaussian kernel. The method assumes overlapping cell populations between batches and that batch effects are orthogonal to biological variation [5].
4. **Scanorama**. Scanorama aligns datasets by identifying transcriptional matches through randomized SVD for dimensionality reduction and approximate nearest neighbor search via Locality Sensitive Hashing. This approach reduces sensitivity to integration order and computational complexity. Validation in the original paper includes t-SNE visualization, silhouette coefficients for clustering quality, and statistical tests to confirm preserved biological signals [13].
5. **scGen**. scGen employs a variational autoencoder (VAE) to encode gene expression into a latent space, using cell type labels to align batches. Biological variations, such as perturbation responses, are modeled through vector arithmetic in this latent space. The VAE’s training implicitly reduces batch effects by prioritizing shared biological features over technical noise. The hyperparameters used for the scGen model include a maximum of 500 epochs, a batch size of 32, early stopping with a patience of 25 epochs, and a dropout rate of 0.2 [12].
6. **ABC (Autoencoder-Based Correction)**. ABC combines a semi-supervised autoencoder with adversarial training. The encoder compresses data into a latent space, while a discriminator adversarially minimizes batch-related signals. Training occurs in three stages: (1) discriminator optimization, (2) adversarial autoencoder training, and (3) joint optimization of reconstruction and cell type classification losses. The architecture includes dropout and ReLU activation for regularization. We trained the models using the default hyperparameters, which include a latent dimension of 64, a dropout rate of 0.1, a reconstruction loss weight of 0.8, an Adam optimizer for all sub-models, and a training duration of 50 epochs [14].
7. **scVI**. scVI is a deep generative model for single-cell RNA-seq data based on variational inference. It formulates gene expression as a hierarchical Bayesian model in which observed counts are generated from a zero-inflated negative binomial distribution, explicitly accounting for technical factors such as sequencing depth and batch effects. Each cell is represented by a low-dimensional latent variable capturing biological variation, while batch annotations are incorporated as covariates in the generative process to disentangle technical from biological sources of variability. Model parameters are learned using stochastic optimization and amortized variational inference, enabling scalable training on large datasets [16]. In our implementation, we trained scVI for 200 epochs (early stopping enabled) on raw counts stored in `adata.layers[“counts”]` with `n_latent=30` and `batch_size=128`. Gene-space batch correction was obtained by decoding normalized expression into a single reference batch (by default, the first batch in sorted order).
8. **ScanVI**. ScanVI extends scVI by adding a semi-supervised objective that leverages cell-type labels through a discrete cell-identity variable, jointly modeling labeled and unlabeled cells while still conditioning on batch to reduce technical variation in the shared representation [17]. In our implementation, we used `scvi.model.SCANVI` with default parameters and obtained gene-space corrected expression by decoding into a reference batch (set to the first batch category).
9. **MIDAS**. MIDAS (Multi-domain Integration for scRNA-seq Data Alignment and Synthesis) is a deep learning framework for single-cell data integration and batch correction. It treats each batch as a separate domain and learns a shared latent representation through an autoencoder architecture trained jointly across domains. Reconstruction and cycle-consistency objectives are used to align batches without requiring cell-type labels. The corrected data are obtained by projecting cells into this common latent space, thereby reducing domain-specific variation while preserving biological structure [18]. In our implementation, MIDAS was trained from scratch for 400 epochs, using the default hyperparameters of the `scmidas` package.

### 3.3 Unsupervised Batch Discrimination Metrics

To quantitatively assess batch correction in an unsupervised manner, we employed several established metrics based on clustering and local neighborhood properties. Specifically, we evaluated the preservation of biological structure and the extent of batch mixing using Adjusted Rand Index (ARI), Average Silhouette Width (ASW), Principal Component Regression (PCR), k-nearest neighbor batch effect test (kBET), and the inverse Local Simpson’s Index (iLISI). For batch-related metrics, calculations were restricted to cells belonging to the set of cell types shared across all batches as in [6].

#### Adjusted Rand Index (ARI)

We computed ARI using the `adjusted rand score` function from scikitlearn. ARI measures the agreement between an unsupervised clustering and a reference annotation. Specifically, we built a *k*-nearest-neighbor (kNN) graph on the first 20 PCA components and obtained cluster assignments using the Leiden algorithm. We then compared the resulting cluster labels to either cell-type annotations or batch labels, yielding two variants: *ARI-Bio Cluster* (cell type vs. clustering) and *ARI-Batch* (batch vs. clustering). When multiple Leiden resolutions were explored, we selected the resolution that maximised the corresponding ARI. Formally, ARI is defined as:

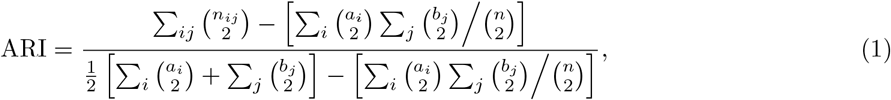

where *n*_*ij*_ is the number of samples shared by cluster *i* in the reference labeling and cluster *j* in the predicted labeling, *a*_*i*_ and *b*_*j*_ are the corresponding cluster sizes, and *n* is the total number of samples. ARI ranges from −1 (anti-agreement) to 1 (perfect agreement), with 0 indicating chance-level agreement.

#### Average Silhouette Width (ASW)

ASW evaluates clustering quality for cell types. The score for each cell *i* is:

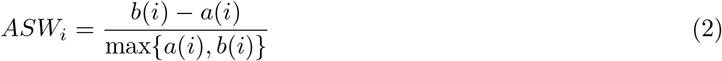

where *a*(*i*) is the average distance between cell *i* and all cells in its cluster, and *b*(*i*) is the average distance to cells in the nearest neighboring cluster. The global ASW is the mean across all cells, ranging from −1 (poor clustering) to 1 (ideal separation). We computed ASW on the first 20 PCA components.

#### Principal Component Regression (PCR)

PCR assesses the extent to which batch information is captured in the principal components of the data. Specifically, it is based on the fitting of a linear regression model to predict batch labels using the first 20 PCA components as input features. The coefficient of determination (*R*^2^) from this regression quantifies the proportion of variance in batch labels explained by the latent space. A high *R*^2^ indicates strong retention of batch effects, while values close to 0 suggest effective batch removal. We report the *R*^2^ averaged across all batches using one-vs-rest encoding.

#### kBET

kBET quantifies local batch mixing by testing whether the batch label distribution in each cell’s neighborhood matches the global distribution. For each cell, a *χ*^2^-test compares its *k*-nearest neighbors’ batch composition to the expected global distribution. Here, we computed kBET on the first 20 principal components using the `scIB` library [46]. The number of nearest neighbors *k* is selected automatically per cluster using the heuristic implemented in `scIB`: it is defined as the minimum value between 70 and one-quarter of the average number of cells per batch within that cluster, ensuring a practical lower bound of 10 neighbors. The final metric is the fraction of cells rejecting the null hypothesis, where 0 indicates perfect mixing and 1 complete batch separation [19].

#### iLISI

iLISI (Local Inverse Simpson’s Index) evaluates batch mixing by measuring the local diversity of batches within the cell neighbors. For each cell *i*, the inverse Simpson’s index is computed over its *k*-nearest neighbors:

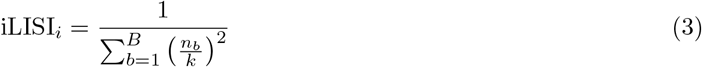

where *n*_*b*_ is the number of cells in batch *b* in the neighborhood of size *k*, and *B* is the total number of batches. Higher values (up to *B*) indicate balanced mixing, while values near 1 suggest batch-specific clustering [11]. The reported value is normalized between 0 and 1, following the implementation used in scIB [46]. iLISI has been calculated on the first 20 PCA components.

### 3.4 Batch Probing Score: a supervised metric to evaluate the batch signal strength after correction

We introduce a supervised approach to directly quantify the residual batch signal after correction. Unlike unsupervised metrics, which rely on geometric properties of the embedding space, this method explicitly measures how much batch information remains accessible to a ML model. Inspired by the ML technique known as probing [27], we define the Batch Probing Score (BPS) as the predictive accuracy of a classifier trained to infer batch labels from the corrected expression data. A high BPS indicates that the batch signal is still learnable, suggesting incomplete correction, while a BPS near zero implies successful batch removal. As with the unsupervised metrics, BPS is computed exclusively on cells belonging to the set of cell types shared across all batches.

#### 3.4.1 Classifiers

First, the expression matrix *X* (rows: cells, columns: genes) is standardized (using the `scikit-learn` *StandardScaler*) to follow a standard normal distribution, i.e., *N* (0, 1).

We perform the probing with two classifiers: a Logistic Regression (LR) and a Random Forest (RF) [47]. For the LR model (L2 regularization, *C* = 1.0), the expression values are further truncated at the 99th percentile to mitigate the influence of outliers (see Suppl. Section S3 for details). The models are trained on *X* and evaluated using the batch labels as the prediction target *y*. The RF classifier is initialized with the following hyperparameters: `n_estimators=200, max depth=50, class_weight=“balanced_subsample”`.

#### 3.4.2 K-Fold cross-validation

To robustly estimate the batch signal available to the model, it is crucial to evaluate its generalization ability. To do so, we use a stratified 5-fold cross-validation (CV). During the CV, the dataset is partitioned into five equal-sized folds while preserving the distribution of the target variable (either batch labels or cell type labels). For each iteration, the classifiers are trained on four folds and tested on the remaining fold, ensuring that each fold serves as the test set exactly once. This protocol minimizes bias due to random data splits and provides an aggregate measure of generalization performance across diverse subsets of the data. The evaluation metrics are reported as the mean value across all folds.

Importantly, in this setting, both training and evaluation of the classifier are performed entirely on the data on which one wishes to quantify the batch signal (i.e., typically the corrected data).

We additionally verified that the computation of BPS is computationally tractable and scales polynomially with the number of cells, with runtime dominated by the cross-validation training of the probing classifiers (see Suppl. Section S5).

#### 3.4.3 Evaluation metrics

To quantify the residual batch signal captured by our BPS, we evaluated two widely used classification metrics: the *Matthews Correlation Coefficient (MCC)* and the *Area Under the Receiver Operating Characteristic Curve (ROC AUC)*.

The *MCC*,

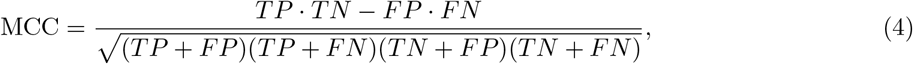

assesses how accurately a classifier assigns each cell to its *exact* batch and ranges from –1 (inverse prediction) to 1 (perfect prediction).

The *ROC AUC*, computed in the multiclass setting with the one-versus-rest (OVR) strategy, instead measures the ability to separate any given batch from the union of all others; a value of 1 denotes perfect separability, while 0.5 corresponds to random guessing, and the metric is independent of the decision threshold.

Although MCC and AUC provide complementary views of performance, OVR AUC is the more sensitive, and therefore more conservative, detector of residual batch signal: it flags a problem as soon as the classifier can distinguish *any* batch from the rest, without requiring exact re-identification. This property becomes increasingly important, as the number of batches increases. Empirically, AUC consistently exceeds MCC across datasets, confirming its stricter nature (see Suppl. Section S2 for details).

We therefore define BPS as a normalized version of the ROC AUC:

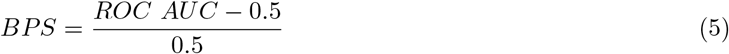

where a BPS close to 0 indicates that there is no detectable batch signal and values approaching 1 suggest strong residual batch effects.

#### 3.4.4 Per-cell-type BPS

To disentangle residual batch signal from cell-type composition imbalance across batches, we additionally compute a per-cell-type variant of BPS-RF. For each cell type *c* present in all batches, a separate RF classifier is trained and evaluated via stratified 5-fold cross-validation on the subset of cells belonging to *c*, using the same hyperparameters described above. The per-cell-type BPS is then defined as the cell-type-frequency-weighted average of the per-stratum OVR AUC scores:

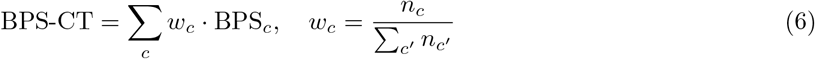

where *n*_*c*_ is the number of cells of type *c* and BPS_*c*_ is the BPS computed on stratum *c*. Because the classifier operates entirely within homogeneous cell-type strata, this variant is blind to composition-driven batch signal by construction and captures only within-cell-type residual batch effects.

### 3.5 Generating a synthetic single-cell dataset

#### 3.5.1 Simulated dataset generation with scsim

We generated *M* = 5 independent in silico single-cell RNA-seq datasets using the `scsim` simulator [48]. Each dataset comprised *G* = 2,000 genes and *N* = 2,000 cells. Cells were grouped into *K* = 10 distinct clusters.

Simulation parameters (e.g. library size distribution, differential-expression probability, dispersion) were set to mimic realistic single-cell experiments, using the default configuration provided in the original `scsim` implementation [48]. For each replicate, raw count matrices and cell metadata (including true cluster/cell type labels) were saved for downstream analysis.

#### 3.5.2 Synthetic batch effect modeling

Let *X* = [*x*_*ij*_] *∈* ℝ^*N ×G*^ denote the unperturbed counts matrix for *N* cells and *G* genes. Each cell *i ∈ {*1, …, *N}* is assigned to a batch *b*_*i*_ *∈ {*0, 1*}*, with half of the cells set to batch 0 and the remaining to batch 1.

To simulate a structured batch effect, a subset of genes is perturbed. For each gene *j, I*_*j*_ ∈ {0, 1} determines whether that gene is affected by the batch perturbation. This activation switch is modeled as a Bernoulli random variable:

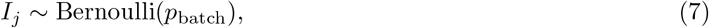

where *p*_batch_ *∈* [0, 1] controls the probability that gene *j* is affected by the batch effect. The perturbed counts matrix *X*^*′*^ = [*x*^*′*^_*ij*_] is then defined as

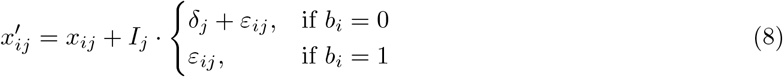

where the batch shift *δ*_*j*_ and noise *ε*_*ij*_ are sampled from

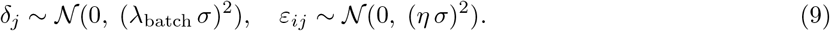

Here, *σ* is the global standard deviation across all entries in the matrix *X*, computed as

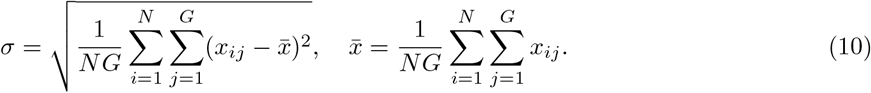

The parameter λ_batch_ controls the magnitude of the batch shift applied deterministically to batch 0 cells only, while *η* regulates the noise intensity applied to all cells. We varied

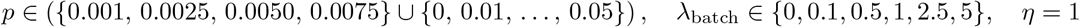

to systematically evaluate the impact of batch perturbations under increasing intensity and sparsity levels. Figure S11 reports the sensitivity scores obtained across all tested combinations of *p* and λ_batch_.

#### 3.5.3 Synthetic biological condition and batch–biology alignment

Starting from the batch-perturbed matrix *X*^*′*^ = [*x*^*′*^_*ij*_] *∈* ℝ^*N ×G*^, we introduce a binary biological condition label *c*_*i*_ *∈ {*0, 1*}* (e.g. case vs control) for each cell *i ∈ {*1, …, *N}*. We set a fixed condition prevalence Pr(*c*_*i*_ = 1) = *π* (here *π* = 0.5).

To explicitly control confounding between batch and biology, we define the batch–biology alignment parameter *α ∈* [0, 1] as a normalized measure of departure from independence between batch and condition:

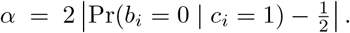

With this definition, *α ≃* 0 indicates that the condition is balanced across batches (no confounding), whereas *α ≃* 1 induces near-complete alignment between condition and batch (maximal confounding). In practice,we sample 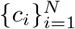 to match the target prevalence *π* and alignment *α* within each dataset.

#### 3.5.4 Biological effect injection

To simulate a structured biological effect, a subset of genes is perturbed. For each gene *j ∈ {*1, …, *G}*, we define a binary indicator 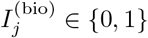determining whether gene *j* is affected by the biological perturbation:

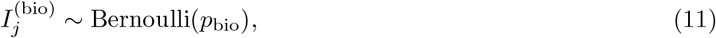

where *p*_bio_ *∈* [0, 1] controls the probability that gene *j* is biologically perturbed.

Starting from *X*^*′*^, we define the biologically perturbed matrix 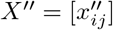 as

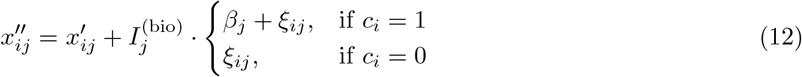

where the biological shift *β*_*j*_ and noise *ξ*_*ij*_ are sampled from

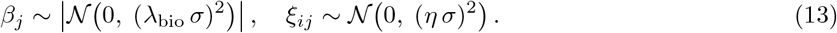

Here, *σ* is the same global standard deviation computed from the unperturbed matrix *X* (as defined above), ensuring batch and biological perturbations are expressed on a comparable scale. *p*_bio_ is set to be equal to 0.1 *\* p*_batch_. The parameter λ_bio_ controls the magnitude of the biological shift applied to condition-positive cells, and it is varied independently from λ_batch_ to decouple biological and technical signal strengths.

#### 3.5.5 Differential-expression recovery

We select differential-expression (DE) analysis as a representative downstream task. For each simulated dataset, we compare cells with *c*_*i*_ = 1 versus *c*_*i*_ = 0 and compute, for each gene *j ∈ {*1, …, *G}*, a two-sided Welch’s *t*-test, obtaining a *p*-value *p*_*j*_.

Genes are ranked by decreasing evidence for DE, using the score *s*_*j*_ = − log_10_(*p*_*j*_). DE recovery is quantified as the area under the precision–recall curve (AUPRC) obtained by treating this ranking as a predictor of ground-truth biologically perturbed genes. Specifically, we define the set of true DE genes as

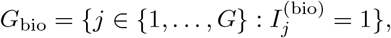

and compute the AUPRC by treating genes in *G*_bio_ as positives and all other genes as negatives, with the rankings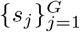.

This yields a direct measure of how well the downstream analysis retrieves the injected biological signal under increasing batch-effect strength and batch-biology alignment.

## Acknowledgments

FC is grateful to all members of the CompBioMed group at the University of Torino for their support and helpful feedback throughout the development of this work. DR is grateful to Anna Laura Mascagni for the constructive discussion.

## Data availability

All datasets used in this study are publicly available. MMCA2 [34, 35], HP5 [36, 37, 38, 39, 40], HPBMC2 [41], and MMHSPC2 [42, 43] were obtained in their preprocessed form from [6]. The HKD11 dataset [44] is available through the Open Problems in Single-Cell Analysis benchmark (openproblems.bio).

## Code availability

The code for this project is available at: https://github.com/codicef/BPS.

## Author contributions

DR and FC conceived the project. FC and DR wrote the manuscript. FC generated the figures and performed the data analysis together with DR. FC, DR, and PF discussed and interpreted the results. DR and PF supervised the study.

