## Supplementary for "On the illusion of scRNA-seq batch effect correction"

#### S1 Comparative evaluation of batch correction: supervised versus unsupervised metrics

Our supervised approach, termed the Batch Probing Score (BPS), is evaluated using a Random Forest (RF) classifier and is based on AUC as the classification metric. The BPS based on RF allows us to sensitively capture non-linear residual batch signals, offering markedly improved detection capabilities over traditional unsupervised metrics such as ARI (Adjusted Rand Index), ASW (Average Silhouette Width), PCR score, iLISI, and kBET. A comprehensive overview of these unsupervised metrics across all datasets and correction methods is shown in the heatmaps in Fig. 1 A-D, Fig. S2 and Fig. S3.

##### S1.1 Sensitivity of BPS versus ARI, ASW and PCR score

As shown in Figures S4 and S5, while most batch correction methods yield near-zero ARI values—suggesting an almost perfect correction—and similarly low ASW scores, these unsupervised metrics fail to capture lingering batch effects. This trend is visually evident in the heatmaps of Fig. 1A (ARI) and Fig. S2 (ASW), where most values sharply drop toward zero after correction. The PCR score (Fig. S6) shows a comparable pattern, and this is further illustrated in panel G of Figure 2, reinforcing the inability of these metrics to detect residual signal.

In contrast, our RF-based BPS consistently reveals significant batch-specific structure that persists after correction. This discrepancy is clearly illustrated in Figure 1G, where ARI, ASW, and PCR estimate an average reduction in batch signals greater than $\sim 85\%$, while Random Forest-based BPS shows only a modest decrease in residual batch signal, averaging around 18%, which indicates that most of the batch effect remains detectable. These results highlight the severe underestimation of residual non-linear batch effects by these unsupervised metrics.

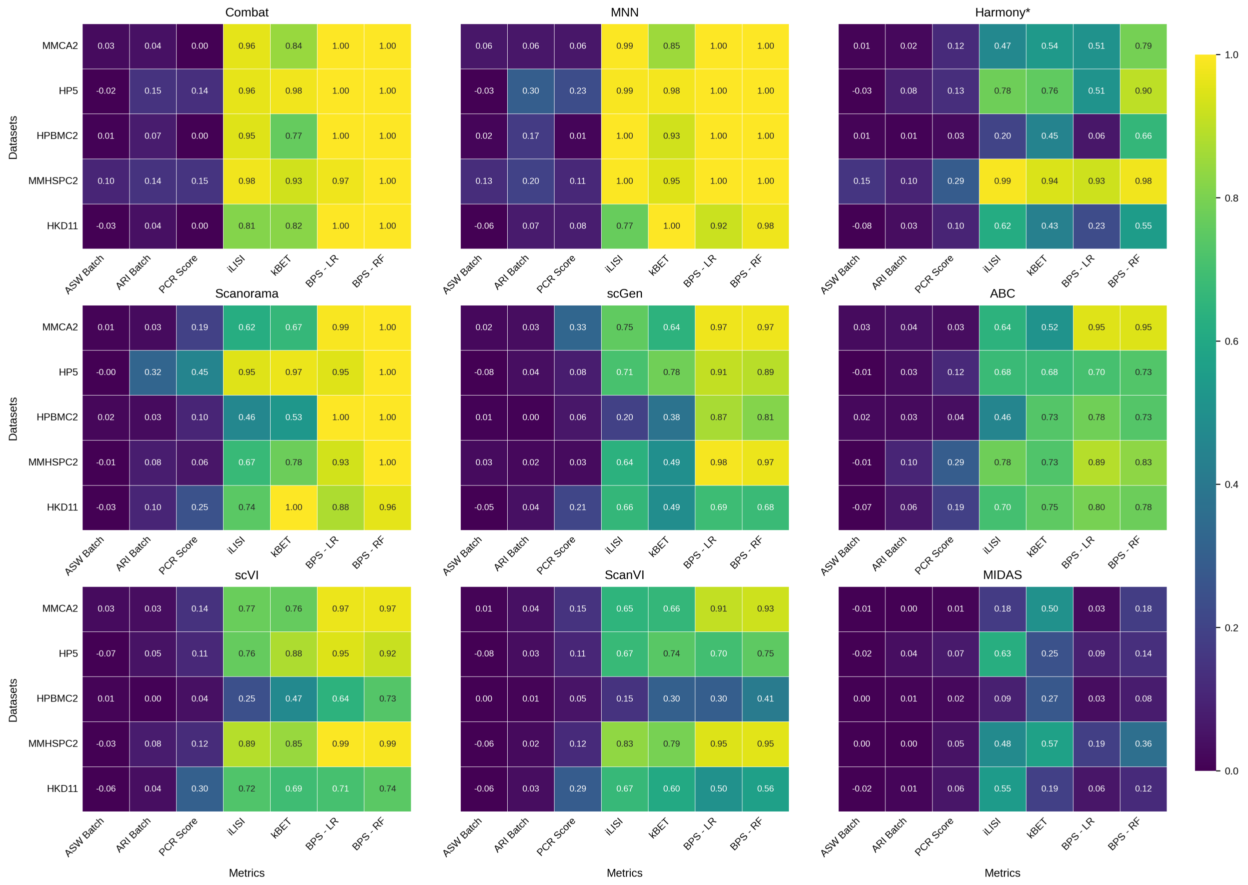

Figure S1: **Multi-panel heatmaps of batch-correction evaluation metrics per method (shared color scale).** Each panel corresponds to one batch-correction method (including Raw non corrected data). Rows denote datasets and columns denote evaluation metrics; values are aggregated across cross-validation folds using the mean. To ensure a consistent direction across metrics, PCR and iLISI are reported as $1 - \text{metric}$ so that larger values indicate *worse* residual batch signal (i.e., poorer batch mixing).

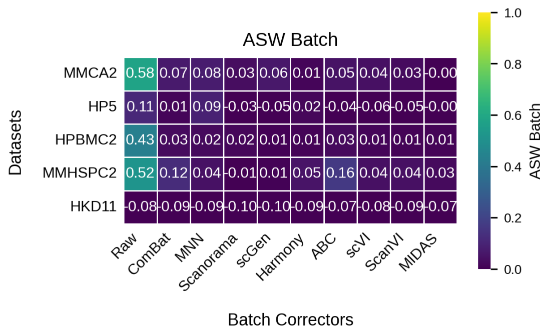

Figure S2: Average Silhouette Width (ASW) Batch correction metric across datasets and methods.

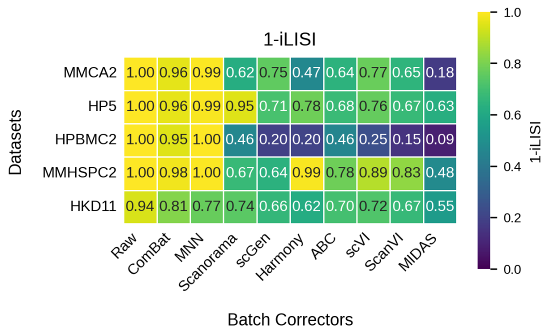


Figure S3: 1- iLISI Batch correction metric across datasets and methods.

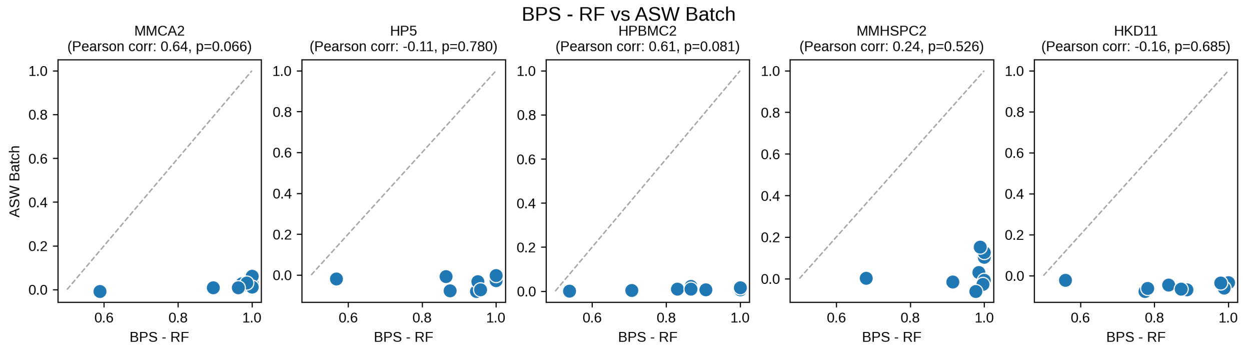


Figure S4: Scatter plots illustrating the relationship between the Batch Probing Score (BPS) based on Random Forest and the Average Silhouette Width (ASW) unsupervised batch correction metric across all datasets. Each point corresponds to the batch correction performance of a different batch correction method, and the Pearson correlation coefficient (r) and p-value are reported for each panel. Raw data have been excluded from the analysis.

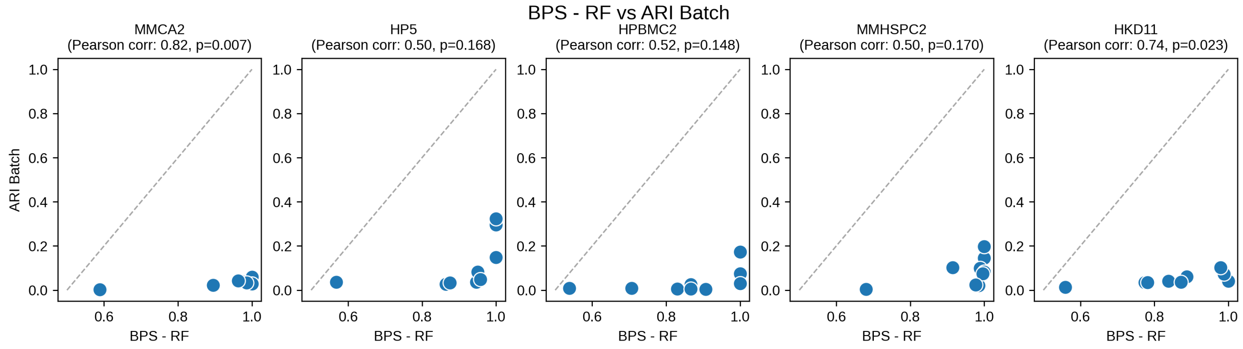


Figure S5: Scatter plots illustrating the relationship between the Batch Probing Score (BPS) based on Random Forest and the Adjusted Rand Index (ARI) unsupervised batch correction metric across all datasets. Each point corresponds to the batch correction performance of a different batch correction method, and the Pearson correlation coefficient (r) and p-value are reported for each panel. Raw data have been excluded from the analysis.

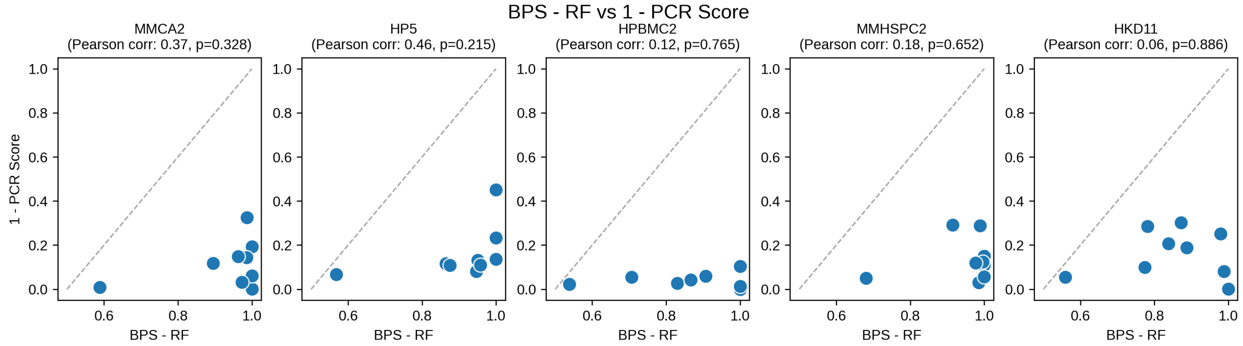


Figure S6: Scatter plots illustrating the relationship between the Batch Probing Score (BPS) based on Random Forest and the Principal Component Regression (PCR) Score unsupervised batch correction metric across all datasets. Each point corresponds to the batch correction performance of a different batch correction method, and the Pearson correlation coefficient (r) and p-value are reported for each panel. Raw data have been excluded from the analysis.

#### S1.2 kBET and iLISI: more sensitive unsupervised metrics

Figure S1 illustrates that unsupervised metrics such as kBET and iLISI provide a less optimistic assessment of residual batch effects compared to the metrics described above. In particular, Figure S7 shows that kBET is able to detect subtle residual batch effects, rarely assigning a ”perfect” score to any correction method. Similarly, Figure S8 indicates that iLISI exhibits comparable behavior. Indeed Figure 1G shows that unsupervised measures such as PCR, ASW, and ARI indicate average reductions in measured batch signal of 86%, 96%, and 81%, respectively. In contrast, both iLISI and kBET estimate much lower average reductions (31% and 29%, respectively), while our supervised BPS-RF reveals an even smaller reduction. This demonstrates that, although unsupervised metrics like kBET and iLISI are more sensitive than ARI or ASW, they still tend to overestimate the efficacy of batch correction. The relatively low correlations between these unsupervised measures and our RF-based BPS (0.66 for iLISI and 0.78 for kBET, as shown in Figure S9) further suggest that they capture complementary, but distinct, aspects of batch effect correction, with our RF-based BPS providing the most sensitive and conservative estimate of residual batch effects.

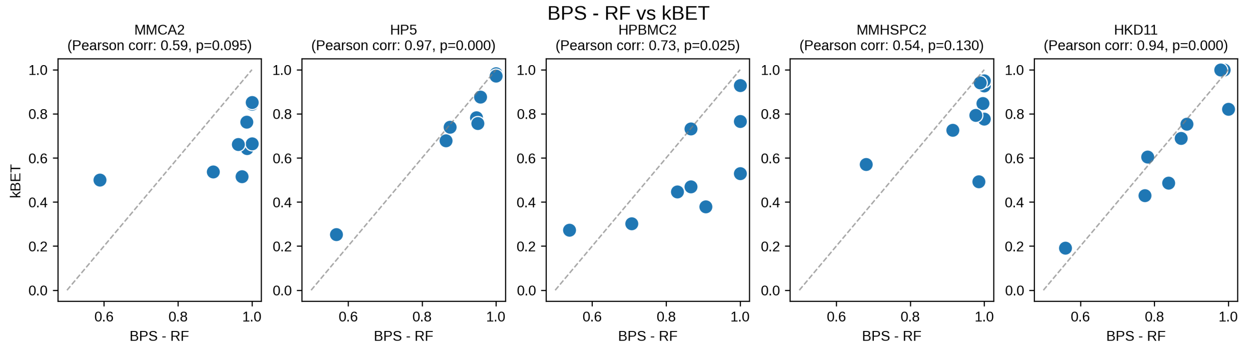


Figure S7: Scatter plots illustrating the relationship between the Batch Probing Score (BPS) based on Random Forest and the kBET unsupervised batch correction metric across all datasets. Each point corresponds to the batch correction performance of a different batch correction method, and the Pearson correlation coefficient (r) and p-value are reported for each panel. Raw data have been excluded from the analysis.

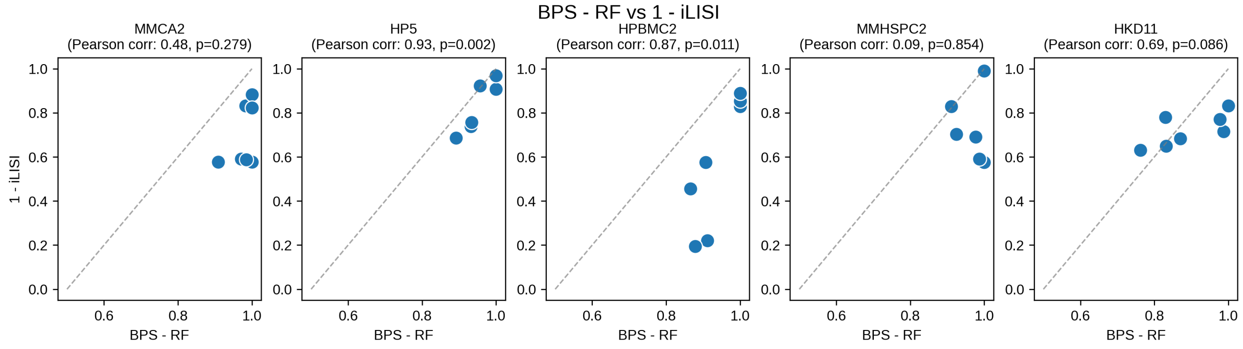


Figure S8: Scatter plots illustrating the relationship between the Batch Probing Score(BPS) based on Random Forest and the 1-iLISI unsupervised batch correction metric across all datasets. Each point corresponds to the batch correction performance of a different batch correction method, and the Pearson correlation coefficient (r) and p-value are reported for each panel. Raw data have been excluded from the analysis.

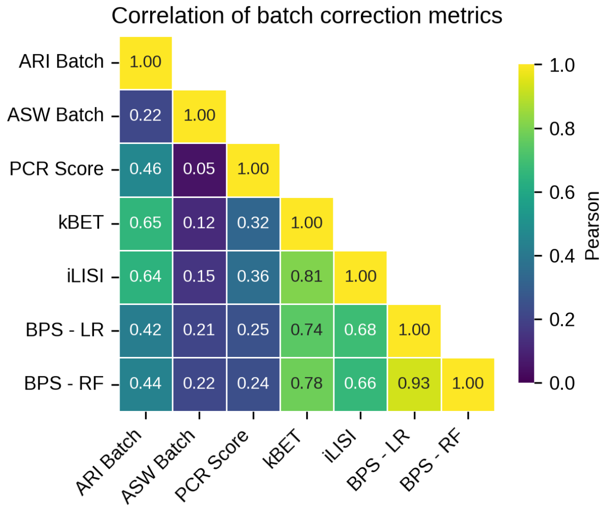


Figure S9: Heatmap showing the Pearson correlation between the different metrics computed across combinations of batch correctors and datasets.

### S2 ROC AUC as a robust and conservative metric of batch correction performance

We defined a Batch Probing Score (BPS) to directly quantify the residual batch signals after correction, using a classification metric to measure batch separability in the corrected data. For this, we considered two candidate metrics: the Matthews correlation coefficient (MCC) and the area under the receiver operating characteristic curve (AUC) (see Methods 3.4). These metrics assess slightly different aspects of batch classifier performance:

- *MCC:* This metric quantifies the classifier's ability to correctly assign each cell **to its respective batch** after correction. Essentially, it evaluates the accuracy with which the classifier identifies **the specific batch** membership of each sample.

- *OVR AUC:* This metric is used in multiclass settings and is computed via the one-versus-rest (OVR) strategy, where the AUC measures the classifier's ability to distinguish one batch from all others. In multi-class problems, it reflects the classifier's ability to **distinguish one batch with respect to the others**, which is more sensitive to subtle residual batch signals than assigning the exact batch of origin, as measured by MCC.

Consequently, OVR AUC can be considered a more conservative measure of residual batch signal, typically yielding higher scores because it is sufficient for the classifier to distinguish between distinct batches. This effect is particularly evident when there is a large number of batches.

In contrast, a high MCC indicates not only that the classifier can differentiate between batches, but also that it can accurately assign each sample to its correct batch, thus representing a more concerning form of residual batch signal.

This discrepancy is evident in Fig. S11, where AUC-based BPS-RF remains high for several dataset-corrector pairs while MCC-based BPS-RF is lower, consistent with OVR AUC capturing batch separability without requiring exact batch assignment.

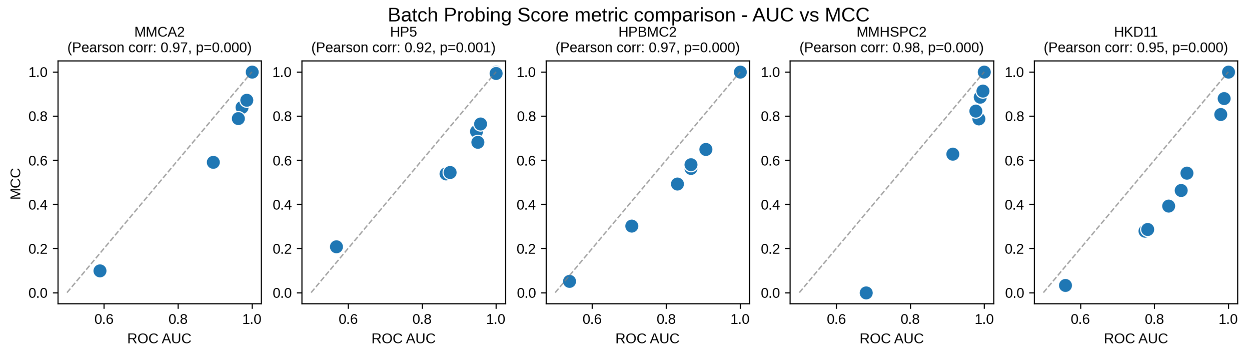


Figure S10: Scatter plots illustrating the relationship between the Matthews correlation coefficient (MCC) and the area under the receiver operating characteristic curve (ROC AUC) across all datasets. Each point corresponds to the classification performance of a different batch correction method, and the Pearson correlation coefficient (r) and p-value are reported for each panel.

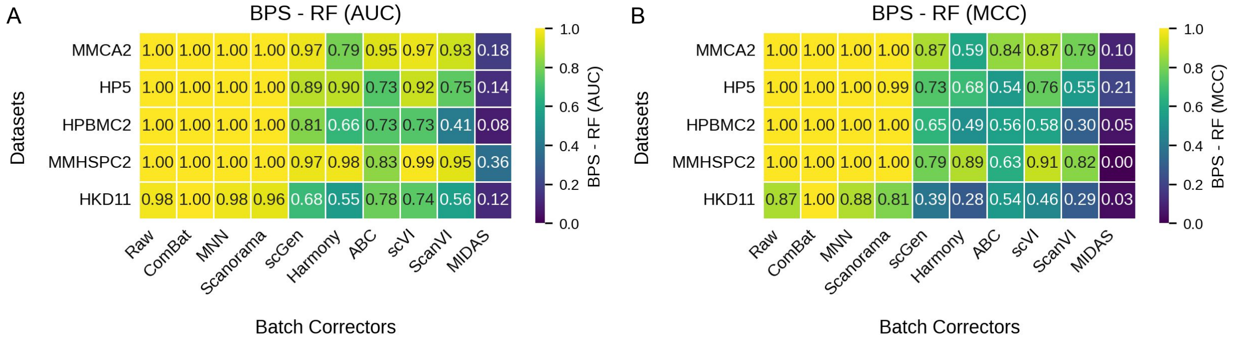


Figure S11: Heatmaps report BPS-RF computed on the same dataset × batch-corrector pairs using (A) OVR ROC AUC and (B) multiclass Matthews correlation coefficient (MCC). Values are shown in each cell and mapped to the same $[0, 1]$ color scale (higher = stronger residual batch separability).

Despite these differences, the two measures are fundamentally related, as evidenced by their strong correlation (see Figure S10). Therefore, we selected ROC AUC as the basis for the BPS due to its conservative nature in detecting residual signals. The BPS is therefore computed using ROC AUC, providing a standardized measure to benchmark batch correction performance across datasets.

### S3 Outlier impact on Logistic Regression prediction

Our analysis of ComBat-corrected expression data revealed an unusual phenomenon: when logistic regression is applied without appropriate preprocessing, the model produces predictions with an AUC below 0.5—indicating inverted (negative) predictions. This reflects a pathological behavior where the model learns a reversed decision boundary due to outlier dominance.

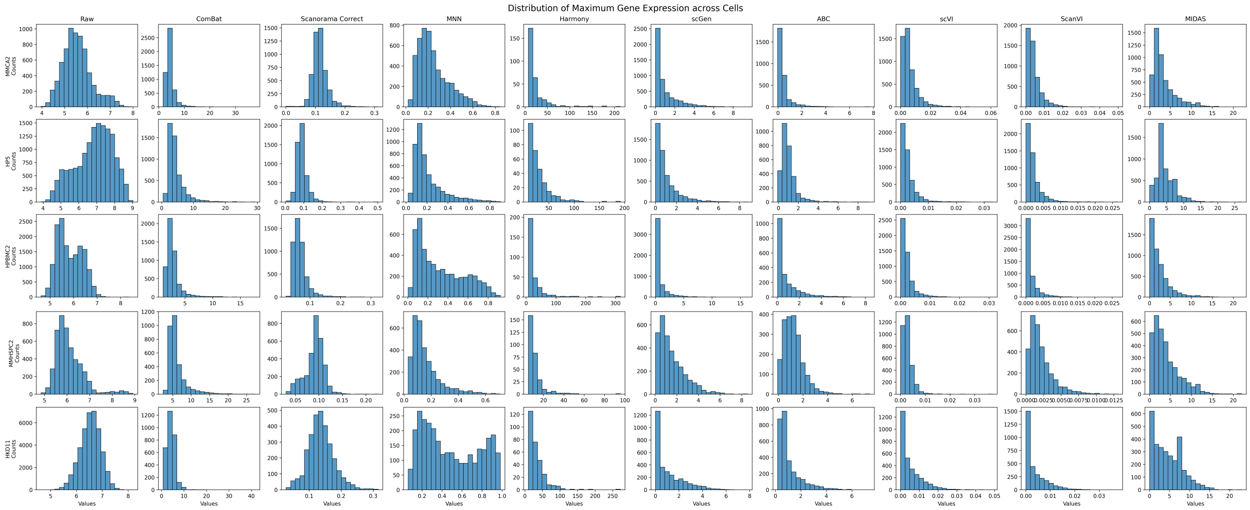


Figure S12: Distribution of maximum gene expression values across genes for each dataset and correction method. Each subplot shows the distribution of the maximum expression value (across all cells) for each gene, for a given dataset (rows) and batch correction method (columns). The first column shows the uncorrected (raw) distributions, while the others reflect the effects of various correction techniques.

Investigation of the expression distributions (Figure S12) demonstrated that they are highly skewed, with an extended upper tail containing extreme values. These outliers heavily distort the training process, leading the logistic regression model to estimate coefficients that drive the predictions in the opposite direction. As illustrated in Figure S13, progressively truncating the extreme values results in a steady improvement in AUC.

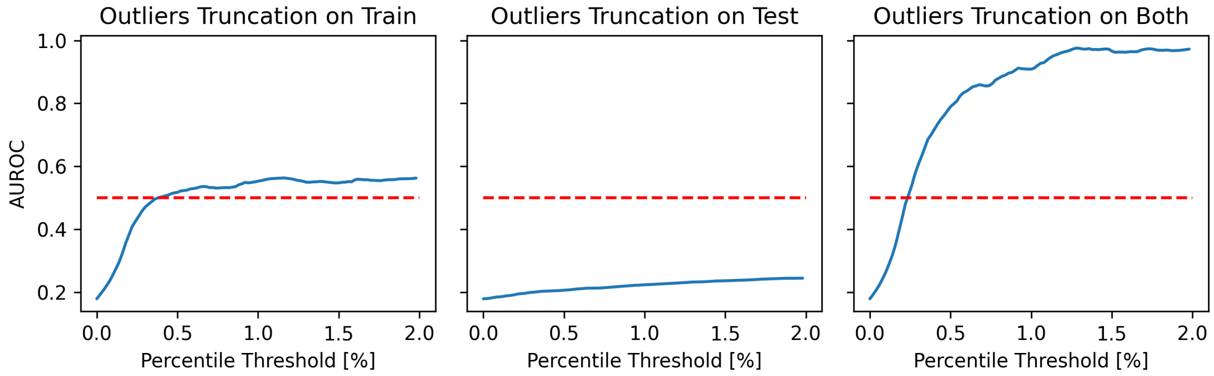


Figure S13: Effect of outlier truncation on logistic regression performance (ROC AUC) in cross-validation on ComBat-corrected MMCA2 dataset. Each panel shows the ROC AUC of logistic regression as a function of the upper and lower percentile threshold used to clip gene expression values, applied respectively to the training set (left), the test set (middle), or both (right). No additional preprocessing was applied beyond the percentile-based truncation. Without truncation, the model exhibits pathological behavior with ROC AUC values below 0.5, indicative of inverted predictions.

Based on these results, we adopted a definitive preprocessing step—truncating the highest 1% of values (capping at the 99th percentile). This intervention stabilizes the estimated coefficients during training, restoring the expected behavior of logistic regression and yielding an AUC close to 1 in many batch correctors-dataset combinations, as demonstrated in main Figure 2D.

In summary, our findings show that unprocessed ComBat data with highly skewed distributions can lead to completely inverted logistic regression predictions, while a simple percentile-based truncation effectively mitigates this issue. Although this phenomenon is most pronounced in ComBat-corrected datasets, similar distortions may occur in other models with extreme values, and in general, truncation leads to a more reliable modeling of the underlying linear signal in the residuals.

### S4 Simulated dataset results

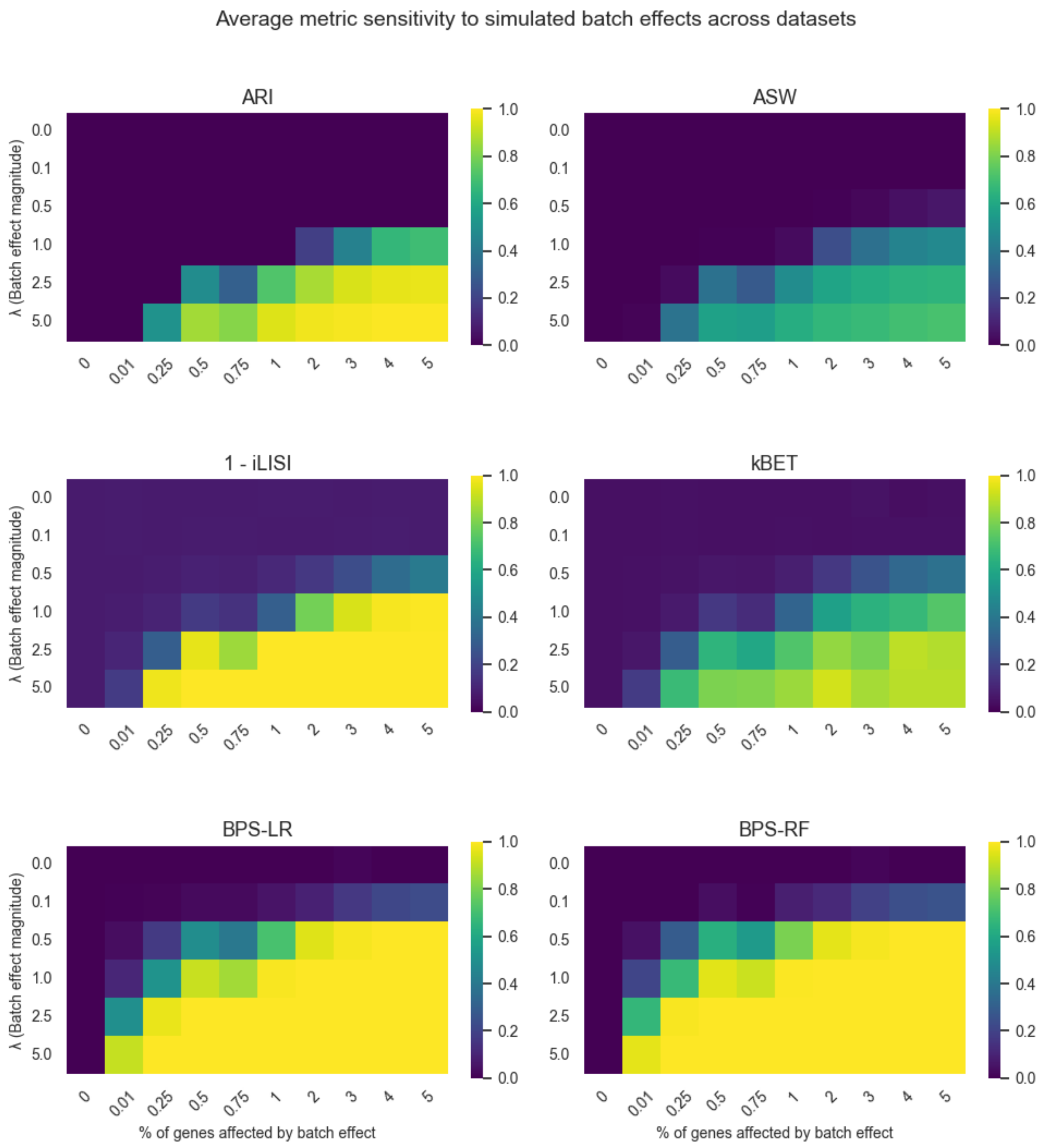


Figure S14: Heatmaps showing the average sensitivity of each metric to simulated batch effects across 5 simulated datasets. For each metric, values are averaged over datasets and plotted as a function of two parameters: the magnitude of the simulated batch effect ($\lambda$, y-axis) and the percentage of perturbed genes (x-axis). Color intensity indicates the metric value (higher implies stronger detection of batch effect), highlighting how each metric responds to increasing batch signal strength and prevalence.

### S5 Runtime and scalability

We empirically evaluated the computational scalability of BPS by measuring runtime as a function of the number of cells on synthetic datasets, keeping the number of genes and batches fixed. BPS consists of two independent probes trained with 5-fold cross-validation: a Random Forest (BPS-RF) and a Logistic Regression (BPS-LR), preceded by feature standardization (and truncation for LR) (see Methods).

Figure S15 shows the runtime scaling of the two probes separately on a log–log scale. As expected, the total cost is dominated by the cross-validation training of the classifiers, while preprocessing (standardization and truncation) contributes only marginally to the overall runtime. BPS-RF is consistently more computationally demanding than BPS-LR, reflecting the higher complexity of the non-linear model.

Importantly, the observed scaling remains approximately polynomial in the number of cells, and BPS can be computed at atlas scale without becoming a practical bottleneck. Since BPS-RF and BPS-LR are independent, they can also be computed separately depending on the desired trade-off between computational cost and sensitivity to non-linear batch signal.

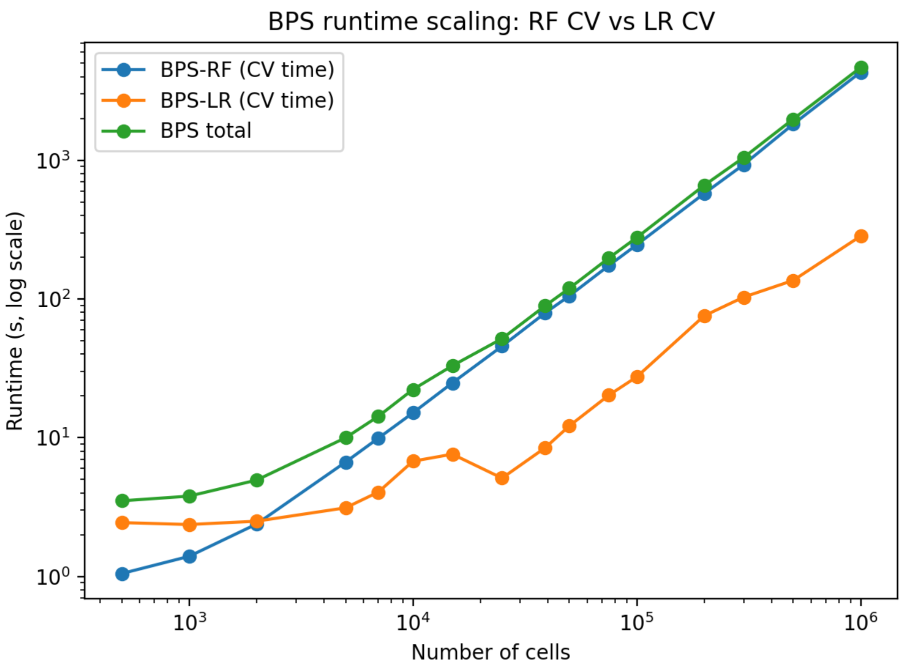


Figure S15: **Runtime scaling of the Batch Probing Score (BPS).** Log–log plot of runtime as a function of the number of cells for the two probing classifiers used in BPS. The Random Forest probe (BPS-RF) and the Logistic Regression probe (BPS-LR) are trained with 5-fold cross-validation. Reported times correspond to the cross-validation phase only. Both components exhibit stable scaling, indicating that BPS can be computed at large scale without becoming a computational bottleneck.

### S6 Cell-type composition per batch across datasets

| Dataset | Cell type | Cells ($n$) | Overall (%) | Batch range (%) |
|---|---|---|---|---|
| *Mus Musculus - Cell Atlas (2 batches)* | | | | |
| | T-cell | 1421 | 20.4 | 16.6–22.9 |
| | Endothelial | 1134 | 16.3 | 10.2–25.9 |
| | B-cell | 1106 | 15.9 | 7.7–28.8 |
| | Stromal | 1032 | 14.8 | 12.2–16.5 |
| | Monocyte | 480 | 6.9 | 3.5–9.1 |
| | Macrophage | 469 | 6.7 | 1.6–10.0 |
| | Neutrophil | 454 | 6.5 | 5.3–7.3 |
| | Epithelial | 390 | 5.6 | 1.3–8.4 |
| | Dendritic | 220 | 3.2 | 0.7–4.8 |
| | Smooth-muscle | 163 | 2.3 | 2.3–2.4 |
| | NK | 85 | 1.2 | 0.8–1.8 |
| *Human - Pancreas (5 batches)* | | | | |
| | alpha | 5100 | 34.5 | 27.1–59.4 |
| | beta | 3826 | 25.9 | 12.7–31.6 |
| | ductal | 1804 | 12.2 | 0.0–21.0 |
| | acinar | 1368 | 9.3 | 0.0–11.2 |
| | delta | 966 | 6.5 | 2.0–9.1 |
| | gamma | 656 | 4.4 | 3.0–9.3 |
| | stellate | 511 | 3.5 | 0.0–5.3 |
| | endothelial | 289 | 2.0 | 0.0–2.9 |
| | mesenchymal | 107 | 0.7 | 0.0–5.9 |
| | macrophage | 55 | 0.4 | 0.0–0.6 |
| | mast | 32 | 0.2 | 0.0–0.3 |
| | epsilon | 28 | 0.2 | 0.0–0.3 |
| | schwann | 13 | 0.1 | 0.0–0.2 |
| | t_cell | 7 | 0.0 | 0.0–0.1 |
| | MHC class II | 5 | 0.0 | 0.0–0.2 |
| *Human - PBMC (2 batches)* | | | | |
| | CD4 T cell | 4450 | 28.8 | 28.0–29.6 |
| | Monocyte_CD14 | 4090 | 26.4 | 23.6–29.5 |
| | CD8 T cell | 3142 | 20.3 | 14.4–25.6 |
| | B cell | 2371 | 15.3 | 14.8–15.9 |
| | NK cell | 593 | 3.8 | 3.7–3.9 |
| | Monocyte_FCGR3A | 561 | 3.6 | 2.5–4.8 |
| | Plasmacytoid dendritic cell | 139 | 0.9 | 0.8–1.0 |
| | Megakaryocyte | 106 | 0.7 | 0.6–0.8 |
| | Hematopoietic stem cell | 24 | 0.2 | 0.1–0.2 |
| *Mus Musculus - HSPC (2 batches)* | | | | |
| | MEP | 1457 | 31.3 | 18.9–40.1 |
| | GMP | 1277 | 27.5 | 6.4–42.3 |
| | CMP | 808 | 17.4 | 17.1–17.6 |
| | MPP | 368 | 7.9 | 0.0–19.2 |
| | LTHSC | 323 | 6.9 | 0.0–16.8 |
| | LMPP | 280 | 6.0 | 0.0–14.6 |
| | Unsorted | 136 | 2.9 | 0.0–7.1 |
| *Human Kidney (11 batches)* | | | | |
| | epithelial cell of proximal tubule | 11478 | 29.3 | 15.7–41.1 |
| | kidney distal convoluted tubule epithelial cell | 8469 | 21.6 | 11.6–32.3 |
| | kidney loop of Henle thick ascending limb epithelial cell | 7749 | 19.8 | 3.0–50.7 |
| | renal principal cell | 3295 | 8.4 | 4.0–15.4 |
| | renal alpha-intercalated cell | 2029 | 5.2 | 2.0–10.0 |
| | endothelial cell | 1702 | 4.3 | 2.4–5.7 |
| | parietal epithelial cell | 1103 | 2.8 | 1.4–5.2 |
| | podocyte | 1069 | 2.7 | 0.9–9.2 |
| | kidney loop of Henle thin ascending limb epithelial cell | 679 | 1.7 | 0.6–8.9 |
| | renal beta-intercalated cell | 670 | 1.7 | 0.4–5.4 |
| | mesangial cell | 385 | 1.0 | 0.5–1.5 |
| | fibroblast | 338 | 0.9 | 0.0–3.3 |
| | leukocyte | 210 | 0.5 | 0.1–4.1 |

Table S1: Cell-type composition of each dataset. *Overall* is the percentage of all cells of that type; *Batch range* is the min–max of the cell type's share across the individual batches (a wide range indicates strong cell-type/batch confounding).

#### S7 BPS specificity under null models and cell-type-stratified evaluation

To empirically assess the specificity of the Batch Probing Score, we evaluated BPS-RF under two null models in which any true batch signal is absent by design, and compared its behavior to that of a stratified variant that preserves cell-type composition.

The **random permutation** null model shuffles batch labels uniformly across all cells, destroying any association between expression profiles and batch of origin. The **stratified (cell-type preserving)** null model permutes batch labels within each cell type separately, ensuring that the cell-type composition of each batch is maintained while still removing the true batch signal.

Importantly, Figure S16 shows that under the stratified null, the global BPS-RF is inflated relative to the random permutation null. This occurs because the global BPS captures any batch-predictable signal, including that driven by cell-type composition imbalance across batches, which is a genuine and measurable component of the batch effect. In contrast, the per-cell-type BPS is evaluated within each cell type independently and is therefore blind to composition-driven signal by construction; accordingly, both null models produce near-zero distributions for the per-cell-type BPS score.

Figure S17 reports BPS-RF computed per cell type and averaged with a cell-type-frequency weighting across datasets and correction methods. This per-cell-type variant isolates genuine within-cell-type residual batch signal from any composition-driven component. The pattern remains broadly consistent with the global BPS-RF reported in the main text (Fig. 2E): classical correctors such as ComBat, MNN, and Scanorama leave substantial batch signal intact within individual cell types, while MIDAS achieves the strongest reduction across all datasets.

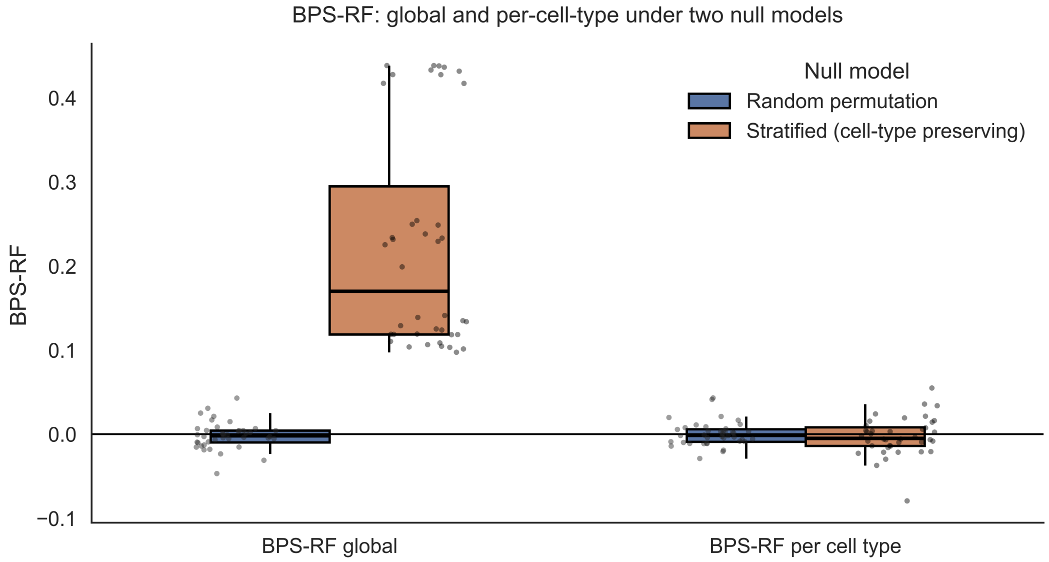


Figure S16: **BPS-RF under two null models.** Distribution of BPS-RF values obtained by permuting batch labels under a random permutation null (blue) and a stratified, cell-type-preserving null (orange), shown separately for the global score and the per-cell-type score. Each point corresponds to one permutation replicate across dataset–corrector pairs.

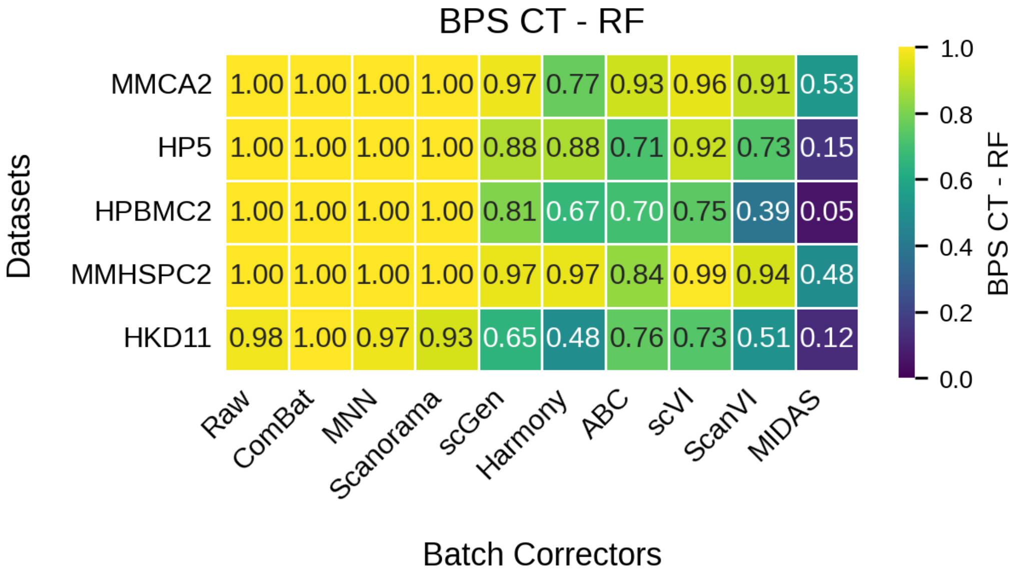


Figure S17: **BPS-RF per cell type (frequency-weighted average) across datasets and correction methods.** Each cell reports the BPS-RF computed within individual cell types and averaged using cell-type frequency as weight, isolating residual batch signal from cell-type composition effects. Rows correspond to datasets and columns to batch correction methods. The color scale follows the same convention as Fig. 2E in the main text (yellow: strong residual batch signal; dark blue/purple: near-complete removal).
